# Immunogenic potential of neopeptides depends on parent protein subcellular location

**DOI:** 10.1101/2021.10.16.464599

**Authors:** Andrea Castro, Saghar Kaabinejadian, William Hildebrand, Maurizio Zanetti, Hannah Carter

## Abstract

Antigen presentation via the major histocompatibility complex (MHC) is essential for anti-tumor immunity, however the rules that determine what tumor-derived peptides will be immunogenic are still incompletely understood. Here we investigate whether protein subcellular location driven constraints on accessibility of peptides to the MHC associate with potential for peptide immunogenicity. Analyzing over 380,000 peptides from studies of MHC presentation and peptide immunogenicity, we find clear spatial biases in both eluted and immunogenic peptides. We find that including parent protein location improves prediction of peptide immunogenicity in multiple datasets. In human immunotherapy cohorts, location was associated with response to a neoantigen vaccine, and immune checkpoint blockade responders generally had a higher burden of neopeptides from accessible locations. We conclude that protein subcellular location adds important information for optimizing immunotherapies.

**Highlights:**

- Peptides eluted from class I and II MHC reflect biases in the subcellular location of the parent proteins
- An embedding-based indicator of parent protein location improves prediction of neoepitope immunogenicity and immunotherapy response
- Neoepitope location improves estimation of effective neoantigen burden and stratification of potential for immunotherapy response

## Introduction

Accurate prediction of immunogenic neopeptides is crucial for the effective application of neoantigen-based cancer treatments such as neoantigen vaccines, immune checkpoint blockade, and adoptive T cell therapy. These immunotherapies all depend on cell surface display of tumor-derived peptides by the major histocompatibility complex (MHC) for immune surveillance by T cells. Current approaches to predict neopeptide immunogenicity (i.e. which tumor mutations will result in neopeptides that are both displayed and recognized by T cells as foreign) largely focus on peptide-MHC affinity, with non-standardized, and sometimes controversial, incorporation of other peptide characteristics such as peptide-MHC stability, agretopicity (Ghorani et al., 2018), foreignness (Łuksza et al., 2017), hydrophobicity, mutation position within the neopeptide (Schmidt et al., 2021), and neopeptide RNA abundance (Wells et al., 2020). However, the utility of these features for predicting immunogenicity varies across experiments and cohorts and ultimately, current tools still yield many false positive neoantigen predictions (Castro et al., 2021; Yadav et al., 2014). Therefore, it follows that there must be other factors that contribute to T cell recognition of peptide-bound MHC that remain to be accounted for.

The canonical pathways by which peptides are added to the MHC for binding differ for class I and class II, with class I peptides requiring transport to the endoplasmic reticulum via TAP transporters (Wieczorek et al., 2017) and class II peptides arriving via endosomes generated through phagocytosis or B cell receptor internalization by antigen presenting cells (Roche and Furuta, 2015). It stands to reason that these distinct pathways could result in peptides from different proteins being more accessible to MHC class I versus class II molecules. Indeed, studies profiling eluted peptide-MHC complexes noted enrichment for peptide origin from intracellular compartments for MHC-I eluted peptides, and for secreted, cell membrane, and extracellular proteins for MHC-II eluted peptides (Abelin et al., 2019). Interestingly, a more recent work found a bias for MHC-I presented peptides to come from proteins with certain molecular functions such as intracellular structural proteins while MHC-II was biased to present membrane transport proteins (Karnaukhov et al., 2021).

These observations imply that proteins in different cellular contexts (location or molecular function, which are correlated (Lu and Hunter, 2004)) can have varying levels of access to the MHC-I or MHC-II presentation pathway. This could further constrain the landscape of peptides that are presented to T cells during cancer development and during T cell selection in the thymus. In the first case, a tumor mutation that would otherwise be effectively bound and displayed by the MHC may not be presented because peptides from the source protein never reach the MHC. For the second, effective presentation of self-antigen during thymic development is required for clonal deletion of corresponding T cells. This pruning of the T cell repertoire is essential to prevent inappropriate activity against self; mutations to the AIRE gene that promotes tissue-specific self-antigen expression during selection result in widespread, multi-organ autoimmunity (Xing and Hogquist, 2012). However, if the peptide repertoire available for T cell selection is constrained by cellular context, it is conceivable that the remaining T cell repertoire would be more capable of mounting a response against peptides from those hidden cellular contexts.

Based on this reasoning, we hypothesized that subcellular location of a source protein could influence the immunogenic potential of its derivative peptides. We collected and analyzed data on peptides presented by the MHC and peptides documented to generate immune responses. To determine the extent to which cellular location can predict neopeptide immunogenicity, we trained and evaluated machine learning models on datasets of experimentally tested neopeptides. FInally, we evaluated source protein subcellular location in the context of immunotherapy response and tumor remodeling by immunotherapy. Together, these analyses support the view that protein subcellular location is a determinant of neopeptide immunogenicity.

## Results

### MHC-bound peptides show enrichment and depletion for certain cellular components

We profiled the subcellular location of peptides bound to MHC class I and class II using gene ontology enrichment analysis (Methods). Peptides eluted from MHC I were obtained from cell lines (Abelin et al., 2017) and 29 normal tissues from the HLA Ligand Atlas (Marcu et al., 2021). Eluted class I peptides were significantly enriched across all samples for 19 cellular components, with the greatest enrichment ratios in cytosol, nucleoplasm, and extracellular exosome components (**Figure 1A**). The integral component of membrane was significantly depleted across all 30 samples, with the integral component of plasma membrane and extracellular region and space being significantly depleted in a majority of class I samples (**Figure 1B**).

**Figure 1.**
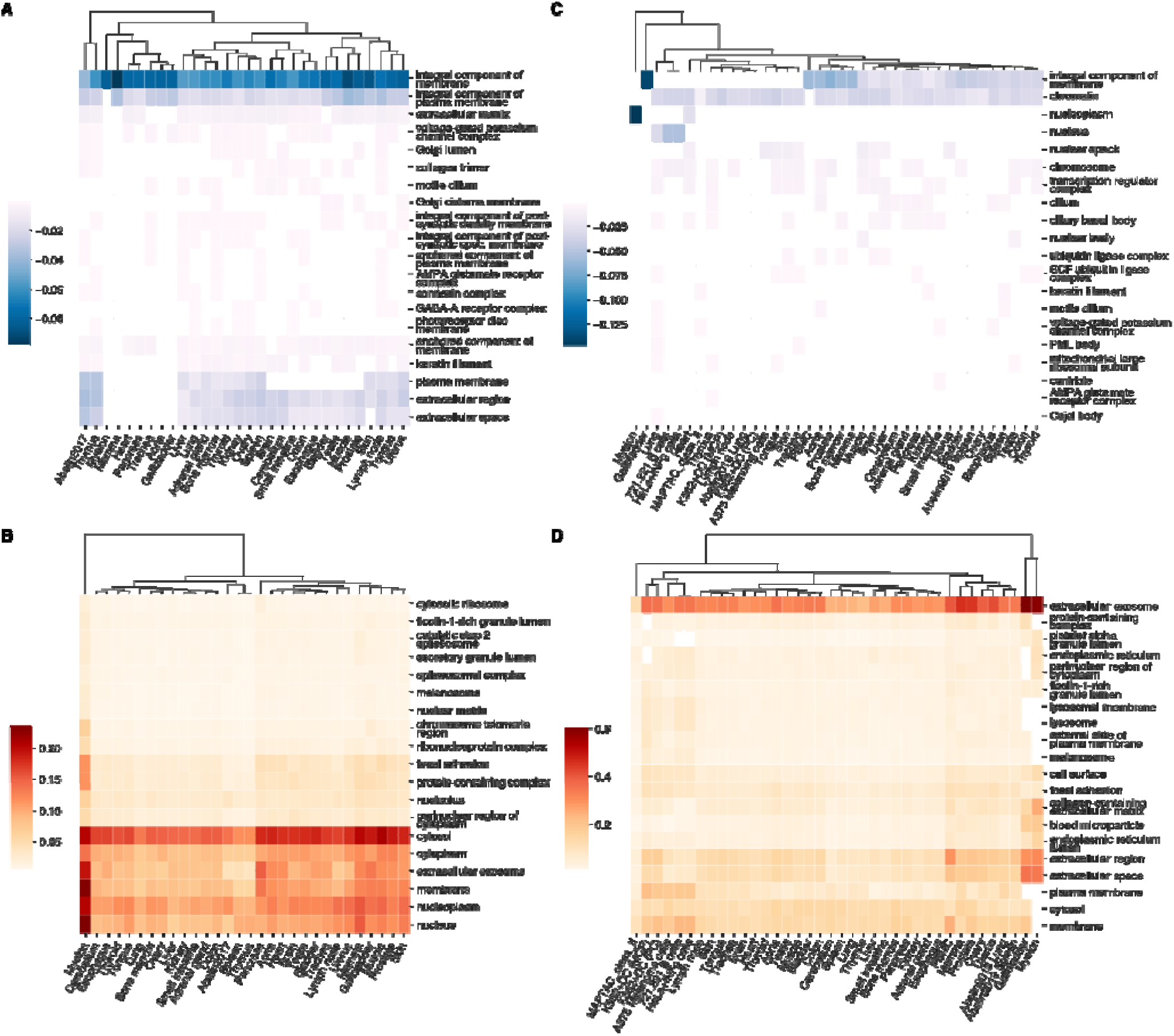
Overview of enrichment or depletion of cellular components in multiple datasets. (A) Clustermap of the top 20 cellular components depleted in eluted peptide-MHC (pMHC) class I from normal and cell lines. (B) Clustermap of 21 cellular components enriched in eluted pMHC-I complexes across all evaluated normal tissues, indicated by “(N)” (Marcu et al., 2021) and evaluated cell lines (Abelin et al., 2017). The color indicates the difference in study vs population enrichment. Clustermaps of (C) depleted and (D) enriched cellular components for eluted pMHC-II from 721.221 B cells, HeLa cells stimulated with IFN-γ, (Abelin et al., 2019), and (Abelin et al., 2019; Marcu et al., 2021).

Peptides bound to MHC II were obtained from 7 different sources including cell lines, normal spleen and lung, and methods of ingestion to APCs (Abelin et al., 2019), 29 normal tissues (Marcu et al., 2021), and our dataset of 721.221 B cells, and IFN-γ treated HeLa cells (Methods). Eluted MHC II peptides were significantly enriched for 35 cellular components across all evaluated samples, with the greatest enrichment of the extracellular exosome, region, and space (top 20 shown in **Figure 1C**). No cellular component was significantly depleted across all evaluated samples; chromatin and the integral component of membrane were significantly depleted in a majority of samples, however there appeared to be more heterogeneity across tissues and cell types (**Figure 1D**).

To better understand why these locations were enriched or depleted for eluted peptides, we compared the expression of genes from enriched or depleted locations using data from the Genotype-Tissue Expression dataset (GTEx) (GTEx Consortium et al., 2017). High gene expression was previously found to correlate with peptide-MHC (pMHC) elution (Abelin et al., 2017). We found that genes from enriched locations had increased expression compared to genes from depleted locations (**Supplementary Figure 1**), though 62% of highly expressed genes were not significantly enriched in eluted pMHC, suggesting that gene expression alone does not drive peptide elution. We also evaluated protein turnover rates from 4 human cell types including B cells, natural killer cells, hepatocytes, and monocytes (Mathieson et al., 2018) as peptides presented by MHC-I are derived from degraded peptides in the cell (Milner et al., 2006). Thus, a high turnover rate could result in more peptides being available for presentation.

Instead, we found that overall, proteins from enriched location categories tended to have longer predicted half lives (**Supplementary Figure 2**), though the cell types evaluated were limited.

These initial findings suggest that proteins in certain subcellular locations may have privileged access to undergo processing in the antigen presentation pathway leading to expression in the context of MHC-I or MHC-II molecules on the cell surface. While presentation by MHC on the cell surface is required for T cell detection, immunogenicity is pre-determined by T cell selection in the thymus during development and depends on self-peptide presentation by medullary thymic epithelial cells (Wang et al., 2020). Analysis of pMHC from 3 normal human thymus tissue samples (Marcu et al., 2021) reflected similar subcellular location biases as other tissues (**Supplementary Figure 3**). Thus, subcellular location is a determinant of the presented peptide landscape across a variety of cellular and tissue contexts, including the thymus where it may influence the composition of the T cell repertoire.

### Certain cellular components are enriched for immunogenic peptides

We hypothesized that the effects of protein subcellular localization bias on peptide availability for presentation and T cell selection would influence the immunogenicity of tumor neoepitopes. Therefore we sought to test whether incorporation of peptide parent protein location would improve prediction of neopeptide immunogenicity. We began by querying the immune epitope database (IEDB) (Vita et al., 2019) for neoepitopes that were assayed for immunogenicity. After filtering (Methods), 2,943 neoepitopes remained, of which 813 (27.6%) were reported to elicit a positive T cell assay result. Cellular component enrichment analysis of parent proteins (n=325) of immunogenic peptides did not reveal any significantly enriched locations, likely due to limited sample size. However, they did tend to come from locations where more eluted peptides were observed on average (**Supplementary Figure 4**). Analysis of unique nonimmunogenic parent proteins (n=772) whose peptides passed the minimum MHC-I binding threshold (<2 netMHCpan percentile rank) showed enrichment for locations including integral component of plasma membrane, plasma membrane, which are depleted for eluted peptide-MHC (**Figure 1B**) but also some other membrane locations that were enriched for MHC-bound peptides, such as the endoplasmic and sarcoplasmic reticulum membranes.

As proteins can localize to multiple locations and have multiple cellular components, we sought a representation capable of capturing complex location patterns. We first used pretrained, 200-dimensional gene ontology embeddings (Kim et al., 2021) to represent each protein’s gene ontology (GO) cellular component annotations, summing embedding vectors if a protein was associated with multiple GO terms. For visualization and use in a machine learning model, we used UMAP dimensionality reduction of these summed embeddings (Methods). As the embeddings and UMAP reduced features are in a non-Euclidean space, proteins with similar locations (e.g. integral component of membrane vs integral component of membrane + plasma membrane) are not necessarily near each other in 2D space (**Supplementary Figure 5**).

We next evaluated locations of immunogenic or non-immunogenic neopeptides from the IEDB, using unique proteins evaluated in respective studies (Methods). As expected, immunogenic and non-immunogenic peptide source proteins had many overlapping locations (**Figure 2A,B**). However, we still observed locations with more immunogenic peptides than not (**Figure 2C**) and vice versa. For example, the mitochondria and mitochondrial matrix had fewer immunogenic peptides while more immunogenic peptides were found in the centrosome and cytoplasmic regions, though small sample sizes and use of several different assays to determine immunogenicity may reduce power.

**Figure 2.**
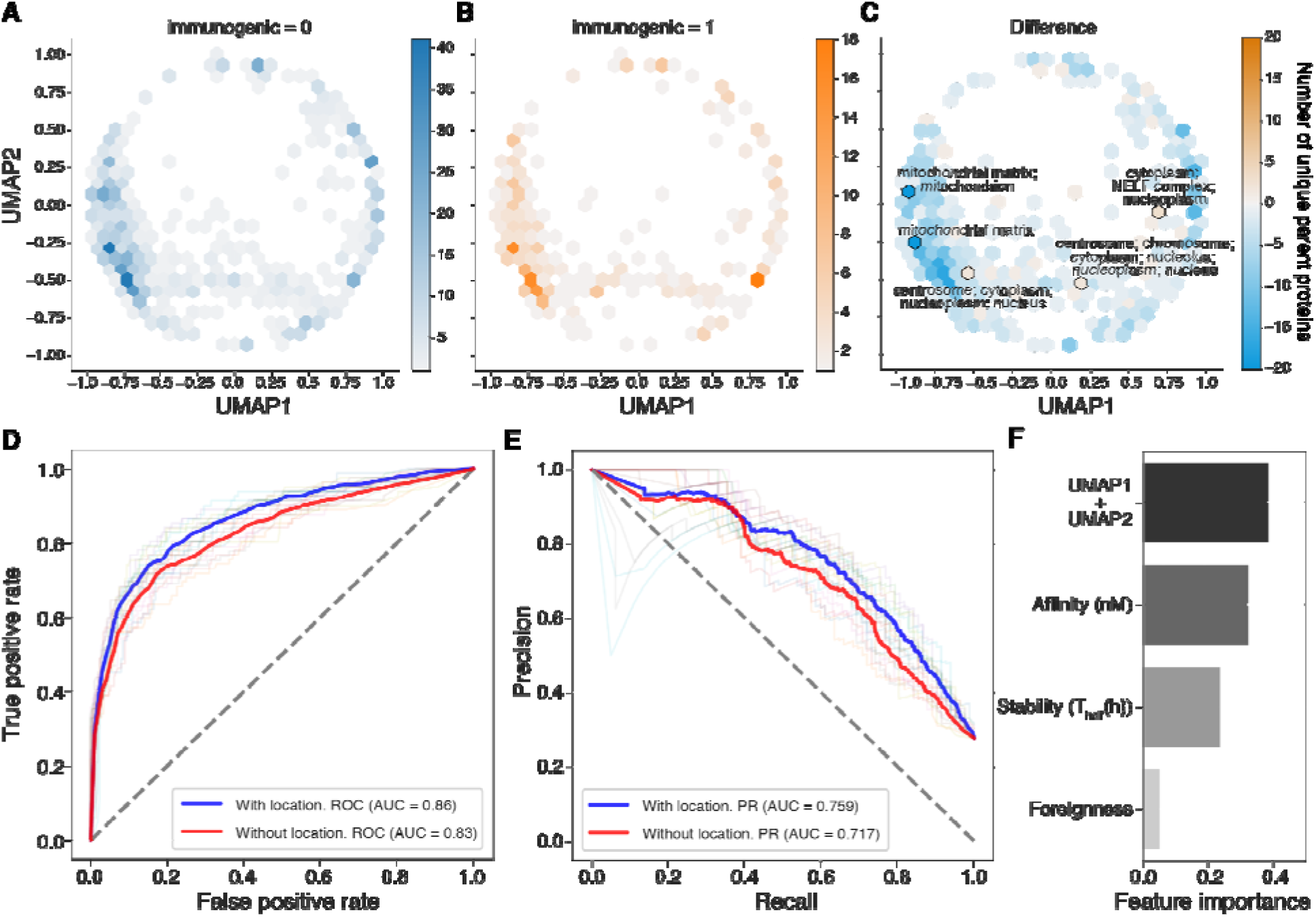
Overview of T cell assayed neoepitopes from IEDB. Hexplots of embedded location for (A) non-immunogenic (blue) and (B) immunogenic (orange) peptides. (C) Hexplot depicting the difference between immunogenic and non-immunogenic hexplots in A and B. Orange indicates more immunogenic peptides, and blue indicates more non-immunogenic peptides. (D) Area under the receiver operating characteristic curve (AUROC) and (E) area under the precision recall curve (AUPRC) for 10-fold cross validation using a Random Forest model incorporating peptide affinity, stability, and foreignness (Methods) with and without parent protein location features. (F) Barplot of model feature importances.

### Parent protein location improves peptide immunogenicity prediction in multiple datasets

Next, we sought to test whether incorporating location as a feature would improve immunogenicity prediction. We performed 10-fold cross validation using a random forest classifier on the IEDB dataset (Methods) with and without adding location to a feature set that comprised peptide-MHC binding affinity (nM) (Jurtz et al., 2017), peptide-MHC stability (Rasmussen et al., 2016), and foreignness (Łuksza et al., 2017; Wells et al., 2020). This dataset did not include many MHC-II peptides, and did not provide enough information about MHC-II alleles to calculate pMHC affinities, therefore we focused on MHC-I peptides. We found that adding location as a feature improved both the area under the receiver operating characteristic (ROC) curve (**Figure 2D**) and precision-recall (PR) curve (**Figure 2E**), and contributed to 38% of the model’s predictive power (**Figure 2F**). We examined the predicted differences between the two models by using the median Youden index to classify peptides as immunogenic or not for each model. 254 peptides were differentially classified between the two models, with 134 now classified as immunogenic and 120 not immunogenic in the location model. The reclassified peptides were enriched for true positives and negatives (Fisher’s exact OR: 1.74, p=0.04), and true positives included peptides from the cytosol while the true negatives included peptides from the nucleus (**Supplementary Figure 6**). Interestingly, among these newly classified peptides, immunogenic versus non-immunogenic peptides had significantly higher median GTEx gene expression but similar affinity, stability, and foreignness scores (**Supplementary Figure 6**), suggesting that location may help predict gene expression. As gene expression was correlated with peptide-MHC elution (**Supplementary Figure 1**) (Abelin et al., 2017), we repeated the analysis, this time including median gene expression obtained from GTEx as a feature. We found that the benefit of including location as a feature persisted, suggesting that location provides distinct information than expression for immunogenicity prediction (**Supplementary Figure 7**).

Next, we tested our model on unseen datasets not included in the IEDB database. First, we analyzed around 900 peptides from Wells *et al*. as this dataset represented the largest collection of immunologically tested peptides we could identify (Wells et al., 2020). As these data were designed to benchmark neoantigen prediction algorithms, they were partitioned into a ~600 peptide discovery set and a ~300 peptide validation set. Initial analysis revealed differences in the distribution of MHC affinity and stability of immunogenic peptides between the IEDB and Wells datasets (**Supplementary Figure 8**) and the parent proteins of immunogenic peptides identified in Wells *et al*., originated from locations that were infrequently observed in the IEDB. Interestingly, while a model trained solely on IEDB did not perform as well as a model trained on the discovery partition provided in the Wells study in ROC analysis (AUROC of 89% vs 93%), it significantly improved the precision-recall curve (AUPRC of 64% vs 9.2%) (**Supplementary Figure 8**). This suggests that filtering candidate neoantigens based on location may significantly reduce false positive predictions. To address systematic differences in the feature sets, we trained a new model combining the IEDB with the Wells discovery set, and were able to achieve a higher recall on the test set (AUROC of 92%) while retaining the benefit for reducing false positives (69% AUPRC) (**Figure 3A-C**).

**Figure 3.**
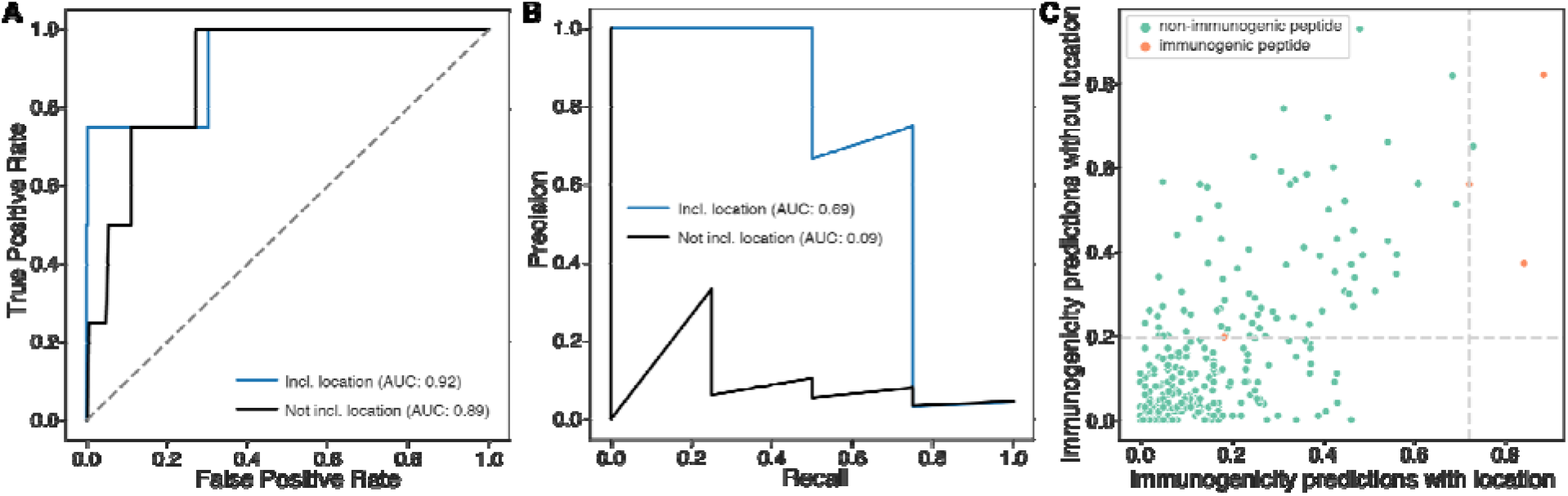
Predicting immunogenicity on unseen datasets. (A) Area under the receiver operating characteristic curve (AUROC) (B) and area under the precision recall curve (AUPRC) for the unseen validation dataset with and without parent protein location features. (C) Scatterplot of the predicted probabilities for unseen test neopeptides to be immunogenic with and without location as a feature. Dashed lines indicate the Youden index for each model, used to optimally threshold predictions. False positives are reduced in the model with location.

Several peptide features including tumor abundance (expression) and agretopicity that were identified in the Wells dataset as being predictive of immunogenicity but were not available in the IEDB dataset. Therefore, we trained a separate model on the Wells discovery set alone incorporating these additional features and tested it on the independent Wells test set. We found that incorporating location improved both the AUROC (89% vs 67%) and AUPRC (9.6% vs 7.5%). In this model, the location features contributed 22% of feature importance, just below affinity (**Supplementary Figure 9**). These experiments suggest that location improves prediction of immunogenic peptides, with the greatest benefit likely coming from reduction in the number of false positive predictions. The large improvement in precision and recall when incorporating the IEDB underscores the benefit of a large training set for capturing the information provided by parent protein location.

We evaluated performance on a second independent dataset of 43 assayed MHC-I neoepitopes from advanced ovarian cancer patients (Liu et al., 2019b) not seen in the IEDB cohort. Of these, 3 (6.9%) were validated as immunogenic. The tested neopeptides once again had significantly different affinity and stability than the IEDB dataset, and while 16 locations were shared between datasets, these did not include the parent proteins for the 3 immunogenic peptides (**Supplementary Figure 10**). We ran the model trained on IEDB alone with and without location features, as well as the model trained on both the IEDB and Wells datasets. We observed improved performance as with the addition of datasets and incorporation of location (**Supplementary Figure 11**), 45% vs 65% AUROC and 6.2% vs 9% AUPRC. Taken together, these findings suggest that immunogenicity prediction benefits from incorporating parent protein subcellular location and can be improved through aggregation of independent datasets across cancer types.

### Immunotherapy response reflects neoepitope parent protein subcellular location

Most T cell based immunotherapies, such as anti-cancer vaccines and immune checkpoint blockade (ICB) depend on the availability of immunogenic peptides to drive effector T cell responses. We speculated that if parent protein subcellular location constrains the set of mutations in a tumor that could potentially be immunogenic, then we should find associations between location and immunotherapy responses. We evaluated three ways in which location might be apparent in human immunotherapy studies. First, we sought to determine whether location was a determinant of T cell response in a neoantigen vaccine study, then we asked whether neopeptides from locations that were more immunogenic were more likely to be depleted by immunotherapy (immunoediting), and finally investigated whether location could improve estimation of the effective neoantigen burden and consequently stratification of responders and nonresponders to ICB.

We first investigated the association of location with immune response in a neoantigen vaccine study (Sahin et al., 2017) and found that parent proteins of neopeptides able to induce a postvaccination response (75/125 tested; 120 distinct parent proteins) were enriched for previously observed immunogenic locations (Fisher’s exact 3.49, p=0.029). Because neoantigen vaccine studies have reported predominantly CD4+ T cell responses (Hilf et al., 2019; Keskin et al., 2019; Ott et al., 2017; Sahin et al., 2017), we also investigated whether the vaccine neopeptides associated with response came from locations from which more MHC-I or MHC-II peptides were eluted by the HLA ligand atlas. The majority of neopeptides tested (92/125, 73.6%) had parent proteins from which peptides had been found eluted from both MHC-I and MHC-II. Nineteen neopeptides’ parent proteins were only observed to be eluted from MHC-I and 7 were exclusive to MHC-II (**Supplementary Figure 12A**). Unlike having parent proteins in a location previously associated with immunogenic peptides, the number of MHC eluted peptides from neopeptide parent proteins alone did not correlate with response, although there was a weak trend for neopeptide parent proteins exclusive to MHC-II to have higher numbers of eluted peptides observed (**Supplementary Figure 12B**). Thus, although we observed correlation between number of eluted peptides and immunogenicity in general (**Supplementary Figure 4**), subcellular location may be a more nuanced determinant of immunogenic potential.

Since we found an association between parent protein subcellular location and post-vaccine response, we speculated that tumor clones cleared during treatment would be more likely to harbor mutations in proteins from immunogenic locations. To further explore this possibility, we evaluated 73 melanoma patients with paired pre- and on-treatment samples to see if there were notable differences between eliminated and persistent neopeptides before and after treatment (Riaz et al., 2017). We focused on responders (n=38, partial/complete response, or >6 months of stable disease) as these patients should have a relatively intact immune response compared to non-responders. While responders had better overall presentation of evaluated neopeptides, eliminated neopeptides did not have significantly better overall MHC allele-specific presentation compared to retained neopeptides in both responders and nonresponders (**Supplementary Figure 13**), suggesting that neopeptide elimination is not driven solely by affinity or stability in this dataset. However, we note that this analysis is complicated by the non-independence of mutations that coexist within the same subclones. To investigate further, we examined 12,915 retained neopeptides from responders that were predicted to be presented by MHC-I (NetMHCpan rank < 2) and, therefore, should have been eliminated by the patient’s immune system. We found that eliminated neopeptides tended to be enriched for locations where immunogenic peptides were previously observed (combining the immunogenic peptides from the IEDB, Wells et al., Liu et al.) (Fisher’s exact OR: 1.08, p=0.09).

Next, we studied the potential for parent protein location to improve ICB response stratification. We evaluated cohorts with whole exome sequencing data, as well as one profiled using a deep sequenced gene panel. We began by looking for association of previously immunogenic locations with response status. We found that in a large cohort of melanoma patients (n=122) (Liu et al., 2019a) where tumor mutation burden associated with response (**Supplementary Figure 14A**), responders compared to nonresponders had a higher burden of proteins from locations where immunogenic responses have been previously observed (Fisher’s exact OR: 1.06, p=0.0014). Considering only mutations from immunogenic locations as putative neoantigens, responders still had a significantly higher burden of neoantigens relative to nonresponders (**Figure 4A**). We repeated this analysis in another melanoma cohort (n=110) (Van Allen et al., 2015), where higher tumor mutation burden (TMB) was also associated with response (**Supplementary Figure 14B**) and this time found no significant association between neopeptide parent protein location and response (Fisher’s exact OR: 0.95, p=0.24). This could be due in part to ignoring other major determinants of immunogenicity such as affinity for the MHC. Therefore, we used a model trained on all datasets with immunogenicity information (IEDB + Wells + Liu (ovarian)) to classify neopeptides in both ICB cohorts as immunogenic or not based on the Youden index of the trained model (Methods). In the Van Allen cohort, the predicted burden of immunogenic peptides in responders was significantly higher than in nonresponders (**Figure 4B**). Comparing predicted neoantigens from models with and without location, to the burden of neopeptides with MHC-I affinity <500nM, we found that filtering out neopeptides predicted not to be immunogenic widened the gap between responders and nonresponders in both cohorts, with the largest difference between responders and non-responders obtained with the model including location (**Figure 4C**), supporting the potential of location to improve stratification of patient groups pre-treatment.

**Figure 4.**
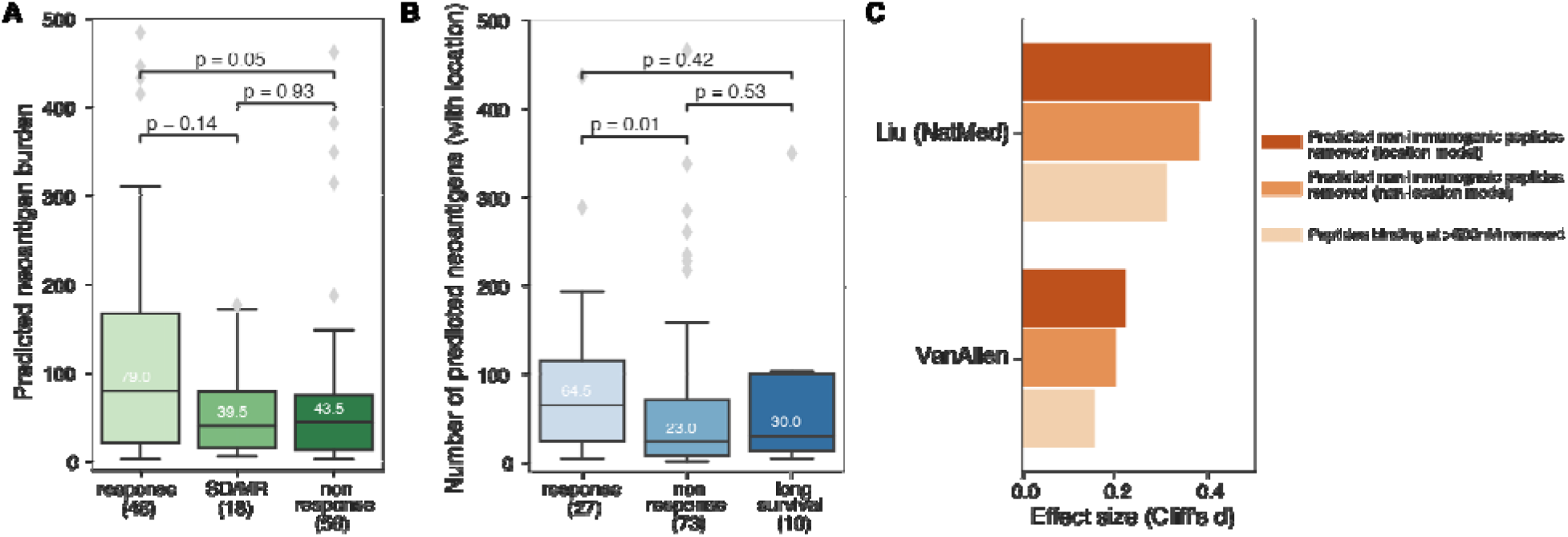
ICB responders carry a higher burden of mutations in proteins from immunogenic locations. A) Predicted neoantigen burden versus response category in the Liu cohort when retaining only mutations in proteins from subcellular locations previously observed to source immunogenic peptides. B) Predicted neoantigen burden versus response category in the VanAllen cohort where neoantigen status is predicted using a model trained on 3 sources of immunogenic peptide and features including peptide MHC affinity, stability, agretopicity and location. C) Barplot of effect sizes between responders and nonresponders where neoantigen status is predicted using a model trained on 3 sources of immunogenic peptide and MHC affinity, stability and agretopicity, with and without location.

Finally, we analyzed a cohort of 83 diverse tumors treated with immune checkpoint monotherapy that were profiled pre-treatment with the Foundation Medicine gene panel. Of 325 genes on this panel, 40 (12.3%) encoded proteins with subcellular locations from which immunogenic peptides had previously been observed, including *ABL1, ALK, APC, ARAF, C11ORF30, EMSY, CCND3, CDK4, CDKN1A, CDKN2A, CREBBP, EGFR, EZH2, FAM46C, FGF19, FGF3, FGF4, FUBP1, GAT A3, ID3, INPP4B, JAK1, KDM5C, KMT2D,MLL2, KRAS, MAP3K1, MDM4, MET, MYCL, MYCL1, NPM1, NT5C2, PALB2, PBRM1, PTPRO, RARA, SMO, TBX3, TET2, TIPARP* and *TP53*. Thirty-seven of these 40 were from regions where only MHC-I peptides had been previously observed, while 2 genes (*BCORL1* and *EPHA3*) were associated with locations from which only MHC-II peptides had been observed.

First, we asked whether the burden of somatic mutations in the 40 genes was informative for stratifying patient outcomes. Focusing on mutation burden in proteins from immunogenic locations reduced the total number of mutations under consideration while preserving the potential to distinguish responders from non-responders and those with stable disease (SD) (**Figure 5A-B**). Second, we asked whether effective presentation of one or more neopeptides from these 40 proteins was a better determinant of outcome than presentation of one or more neopeptides across all proteins in the panel. For this analysis, we focused on the 71 out of 83 patients that carried at least one mutation in these 40 genes. While patient MHC genotype specific presentation scores (PHBR scores, (Marty et al., 2017)) were able to stratify responders from non-responders when all proteins were considered (**Figure 5C**), the stratification improved when we focused on only the 40 proteins from immunogenic locations (**Figure 5D**). We revisited this analysis using a Cox Proportional Hazards model with covariates as described previously (Goodman et al., 2020), and found that when we focused on the 40 panel genes encoding proteins from immunogenic locations, the PHBR score was more significantly associated with outcome (**Supplementary Table 1**). Altogether these results support that subcellular location of parent proteins is a determinant of the effective neoantigen burden in the setting of immunotherapy.

**Figure 5.**
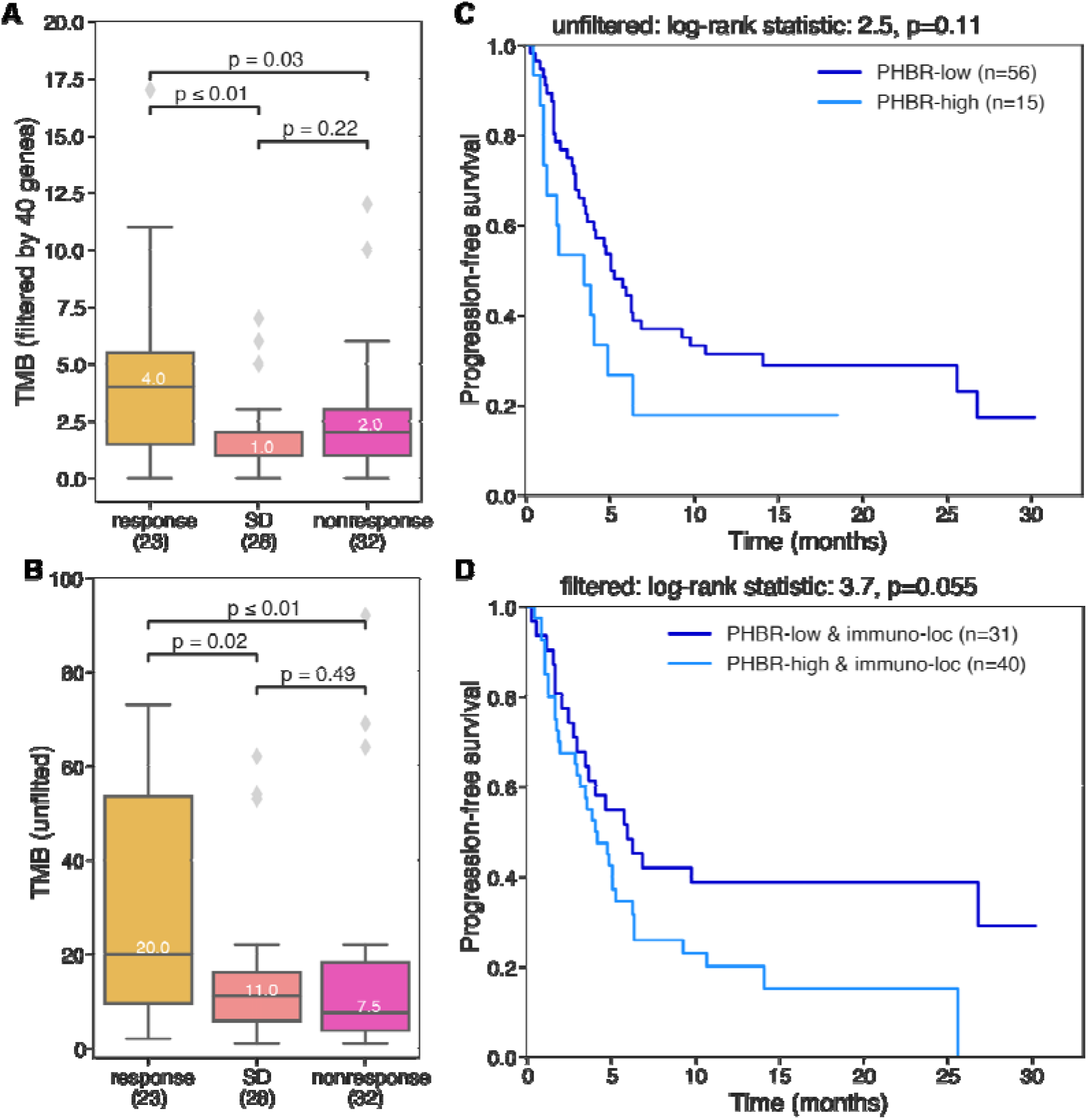
Focusing on immunogenic locations improves response prediction in a gene panel profiled cohort. Tumor mutation burden (A) focusing on the 40 genes whose proteins localize to previously observed immunogenic subcellular locations and (B) all genes in the gene panel. Kaplan Meier curves showing the effect of the best presented mutation on progression-free survival (C) using all genes in the panel and (D) using only the 40 genes of interest.

## Discussion/Conclusion

While immunotherapy has generated more durable responses than targeted therapies (Ledford, 2016), the fraction of patients that respond is lower. Notably, immunotherapy tends to have higher response rates in tumor types with a high burden of somatic mutations which is thought to be a proxy for having a large number of immunogenic mutations. Mapping the mutations in a tumor genome to the subset that are likely to create immunogenic neoantigens is therefore important for realistically assessing the potential for immunotherapy response as well as for designing effective cancer vaccines. Consequently, a variety of metrics have been developed to reveal putative neoantigens in tumor genomes, with the most common being peptide-MHC binding affinity, peptide-MHC complex stability, peptide agretopicity, foreignness, and mutation expression. Here we analyzed peptides from eluted peptide-MHC and found that the subcellular location of proteins also influences which peptides are presented by the MHC. Using a highdimensional cellular location embedding that captured multi-localization mapped to a 2 dimensional representation, we analyzed the implications of parent protein location relative to peptide immunogenicity and immunotherapy response. Immunogenic peptides were biased toward specific subcellular locations and a higher burden of mutations from these regions was associated with more benefit from immunotherapy in multiple cohorts. These findings provide the first evidence that parent protein locations influence both neopeptide presentation and T cell recognition and elimination.

We evaluated both the subcellular locations of proteins from which MHC-I and MHC-II bound peptides originate as well as those associated with peptides labeled as immunogenic based on experimental assays. We note that these locations may not fully overlap. While stable presentation by the MHC is a prerequisite for immunogenicity, it’s possible that not all locations from which peptides are sourced will generate immunogenic peptides. This is dependent in part on the extent of thymic selection. Furthermore, we note that not all experimental assays used to profile immunogenicity will fully recapitulate the dependence on protein location, which could lead to the appearance of some immunogenic peptides coming from regions with no HLA presented peptides. Furthermore, there was substantially less information about MHC-II peptides than MHC-I, leading to more limited assessment of immunogenicity in locations where peptides were predominantly eluted from MHC-II. Nonetheless, more MHC-II presented peptides than MHC-I presented peptides were associated with vaccine response in multiple vaccine studies (Hilf et al., 2019; Keskin et al., 2019; Ott et al., 2017; Sahin et al., 2017).

In general, we speculate that the location constraint could more strongly affect peptide availability for MHC-I. Peptides from different compartments within the cell may have more variable access to the ER, which depends largely on transport from the cytoplasm by TAP family transporters (Yewdell and Bennink, 1999). Peptides displayed by MHC-II come mostly from proteins internalized by antigen presenting cells, however MHC-I and MHC-II has been found abundantly in extracellular exosomes derived from B cells, which may explain the significant enrichment in eluted peptides (Colombo et al., 2014; Wubbolts et al., 2003). In addition, the diversity and availability of such proteins could change drastically in the presence of apoptotic or necrotic cells in the tumor immune microenvironment, making proteins from previously unavailable locations more accessible. Cross-priming may allow some exceptions to location constraints as well (Kurts et al., 2010).

These considerations are particularly important in the context of cancer vaccines. Effective vaccine design depends on selecting peptides that will induce robust immune responses. Inclusion of peptides that stimulate T cell expansion but are not effectively displayed by the MHC at the tumor site creates the risk of generating immunodominance toward ineffective targets (Yewdell and Bennink, 1999). The resulting T cell expansions could be dominated by clones incapable of suppressing the tumor, while more relevant clones are outcompeted in competition for antigen on the APCs (Garcia et al., 2007), nutrient starved (Kedia-Mehta and Finlay, 2019) and may become more easily exhausted (Malandro et al., 2016). Thus, it may be important to avoid including peptides from parent proteins that are less accessible to the MHC. More stringent constraints on peptide accessibility to MHC-I might make selection of effective peptides for MHC-I more challenging than for MHC-II.

Another possible consideration is whether biases in protein location during thymic tolerance render the T cell repertoire more sensitive to proteins from certain locations. Including peptides from these locations could be beneficial. This also leads to the speculation that protein localization changes in tumor cells could alter accessibility to the MHC. If these proteins were less subject to thymic tolerance, they could potentially be more potently immunogenic. One study found that an inverted form of melanoma antigen with altered localization, Melan-A, was recognized by T cells while the native orientation and a variant expressed in the cytosol were not (Rimoldi et al., 2001). Although alterations in localization signals are reportedly rare (Laurila and Vihinen, 2009; Wang and Li, 2014), differences in trafficking could be more common (Tzeng and Wang, 2016). For example, we found that mitochondrial regions were depleted for immunogenic proteins, however a mitochondrial derived vesicles may provide a pathway for proteins from this region to the MHC (Matheoud et al., 2016).

We note several limitations to our study. The pretrained location embeddings were based on characteristics of normal cells, and will reflect any biases or gaps present in the Gene Ontology (Gaudet and Dessimoz, 2017). Furthermore, many proteins map to multiple locations (Thul et al., 2017) and have multiple associated cellular component terms. In this study we weighted each component equally, but it is likely that some locations may be predominant or transient. Immunogenicity is based on experimental assays in the IEDB performed on 325 proteins by various groups using various assays. These proteins could reflect selection bias. Similarly, locations associated with MHC eluted peptides may reflect the specific alleles that were profiled. In addition, MHC II datasets may be biased toward B cells, whereas differences in internalization mechanisms among antigen presenting cell types such as dendritic cells or macrophages could create differences in which proteins are more accessible.

Despite these limitations, we found that incorporating protein location into analysis of immunotherapy cohorts was helpful in several ways. We used location to revise the effective neoantigen burden in tumors and better stratify potential for immunotherapy response. While expression and location provided independent benefit for inferring immunogenicity, correlation between these measures suggests that location could serve as a generic proxy for expression in studies where expression was not directly measured. More insight may be gained from future single cell studies where it is possible to define the clonal architecture of tumors and cosegregating mutations. Location information was also beneficial in a cohort that was profiled with a gene panel, suggesting that this information could be relevant for data more commonly generated in clinic settings. Thus, we conclude that protein subcellular location contributes to shaping the tumor-immune interface and can potentially be leveraged to improve the effective application of immunotherapies.

## Acknowledgements

This work was supported by Emerging Leader Award from The Mark Foundation for Cancer Research grant #18-022-ELA and NIH U24CA248138-01A1 grant subaward 20051-01-144-384 to HC and NIH grant R01CA220009 to HC and MZ. Some analyses herein used data from the GTEx project, which was supported by the Common Fund of the Office of the Director of the National Institutes of Health, and by NCI, NHGRI, NHLBI, NIDA, NIMH, and NINDS.

## Author Contributions

Peptide elution experiments were performed by SK under supervision of WH. Data analysis was performed by AC. Manuscript was written by AC, HC, MZ.

## Data and Methods

### GO analysis

Gene ontology enrichment analysis was performed using GOATOOLS (https://github.com/tanghaibao/goatools) (Klopfenstein et al., 2018) using the standard parameters, and retaining enriched or depleted results if the Benjamini-Hochberg corrected p-value was less than 0.05.

### Abelin 2019 peptides

Peptides were obtained from the published Supplementary Data S1B. Peptides were mapped to parent UniProt sequences and filtered out if they mapped to multiple parent proteins. 69653/76561 (90.7%) peptides uniquely mapped to 1 parent protein sequence.

### Isolation and purification of HLA-DR bound peptides

The human B cell lymphoblastoid cell line 721.221 was grown in complete RPMI 1640 medium (Gibco) supplemented with 10% fetal bovine serum (FBS; Gibco/Invitrogen Corp). The HeLa cell line was grown in DMEM/F12K (Gibco) supplemented with 10% fetal bovine serum (FBS; Gibco/Invitrogen Corp). The cells were grown in large scale cultures in roller bottles and the cell viability was maintained at >90% throughout the experiments. In order to induce HLA-Class II surface expression, HeLa cells were treated with IFN□ (500 U/mL) for 72hrs after which the cells were harvested, washed twice with ice cold PBS and spun down at 2500xg at 4C for 10 minutes. The cell pellets were snap frozen in LN_2_ and stored at −80 until downstream processing. Both cell lines were subjected to high-resolution sequence-based HLA typing (HLA-A, -B, -C, DR, DP and DQ) for authentication prior to large scale culture and data collection.

HLA-DR molecules were purified from the cells by affinity chromatography using the anti-human HLA-DR antibody (clone L243) coupled to CNBr-activated Sepharose 4 Fast Flow (Amersham Pharmacia Biotech, Orsay, France) as described previously with some modifications (Purcell et al., 2019). Briefly, frozen cell pellets were pulverized using Retsch Mixer Mill MM400, resuspended in lysis buffer comprised of Tris pH 8.0 (50 mM), Igepal, 0.5%, NaCl (150 mM) and complete protease inhibitor cocktail (Roche, Mannheim, Germany). Lysates were centrifuged at 200,000 xg for 90 min in an Optima XPN-80 ultracentrifuge (Beckman Coulter, IN, USA) and filtered supernatants were loaded on immunoaffinity columns. After a minimum of 3 passages, columns were washed sequentially with a series of wash buffers (Purcell et al., 2019) and were eluted with 0.2 N acetic acid. The HLA was denatured, and the peptides were isolated by adding glacial acetic acid (up to 10%) and heat. The mixture of peptides and HLA-DR was subjected to reverse phase high performance liquid chromatography (RP-HPLC).

### LC-MS/MS Analysis

Reverse-phase high performance liquid chromatography (RP-HPLC) was used to reduce the complexity of the peptide mixture eluted from the affinity column. First, the eluate was dried under vacuum by using a CentriVap concentrator (Labconco, Kansas City, Missouri, USA). The solid residue was dissolved in 10% acetic acid in water and fractionated over a 150-mm long Gemini C18 column, pore size 110 Å, particle size 5μ (Phenomenex, Torrance, California, USA) by using a Paradigm MG4 instrument (Michrom BioResources, Auburn, California, USA). An acetonitrile (ACN) gradient was run at pH 2 using a two-solvent system. Solvent A contained 2% ACN in water, and solvent B contained 5% water in ACN. Both solvent A and Solvent B contained 0.1% trifluoroacetic acid (TFA). The column was pre-equilibrated at 2% solvent B. After loading the peptide mixture on the column in a period of 19 min by using a solvent system comprised of 2% Solvent B with the flow rate of 160 μL/min, two linear gradients were run at the same flow rate: 4% to 40% Solvent B for 40 min, followed by 40% to 80% Solvent B for 8 min. The percentage of Solvent B was maintained at 80% for 4 min, and then decreased to 2% over a period of 3 min. Fractions were collected in 2 min intervals using a Gilson FC 203B fraction collector (Gilson, Middleton, Wisconsin, USA), and the ultra-violet (UV) absorption profile of the eluate was recorded at 215 nm wavelength.

Peptide-containing HPLC fractions were dried, resuspended in an aqueous solvent composed of 10% acetic acid, 2% ACN and iRT peptides (Biognosys, Schlieren, Switzerland) as internal standards. Fractions were applied individually to an Eksigent nanoLC 415 nanoscale RP-HPLC (AB Sciex, Framingham, Massachusetts, USA), including a 0.5-mm long, 350 μm internal diameter Chrom XP C18 trap column with 3 μm particles and 120 Å pores, and a 15-cm-long ChromXP C18 separation column (75-mm internal diameter) packed with the same medium (AB Sciex, Framingham, Massachusetts, USA). An ACN gradient was run at pH 2.5 using a two-solvent system. Solvent A was 0.1% formic acid in water, and solvent B was 0.1% formic acid in 95% ACN in water. The column was pre-equilibrated at 2% solvent B. Samples were loaded at 5 μL/min flow rate onto the trap column and run through the separation column at 300 nL/min with two linear gradients: 10% to 40% B for 70 minutes, followed by 40% to 80% B for 7 minutes.

The column effluent was ionized using the nanospray III ion source of an AB Sciex TripleTOF 5600 quadrupole time-of-flight mass spectrometer (AB Sciex, Framingham, MA, USA) with the source voltage set to 2,400 V. Information-dependent analysis (IDA) of peptide ions was acquired based on a survey scan in the TOF-MS positive-ion mode over a range of 300 to 1,250 m/z for 0.25 seconds. Following each survey scan, up to 22 ions with a charge state of 2 to 5 and intensity of at least 200 counts per second were subjected to collision-induced dissociation (CID) for tandem MS analysis (MS/MS) over a maximum period of 3.3 seconds. Selection of a particular ion m/z was excluded for 30 seconds after three initial MS/MS experiments. Dynamic collision energy was utilized to automatically adjust the collision voltage based upon ion size and charge. PeakView Software version 1.2.0.3 (AB Sciex, Framingham, MA, USA) was used for data visualization.

### Peptide Identification and Source Protein Information

Peptide sequences were assigned to resulting fragment spectra using PEAKS Studio 10 software (Bioinformatics Solutions, Waterloo, Canada) at a precursor mass error tolerance of 30 ppm and a fragment mass error tolerance of 0.02 Da. A database composed of SwissProt Homo sapiens (taxon identifier 9606) was used as the reference for database search. Variable post-translational modifications (PTM) including acetylation, deamination, pyroglutamate formation, oxidation and sodium adducts were included in database search. Identified peptides were further filtered at a false discovery rate (FDR) of 1% using PEAKS decoy-fusion algorithm. UniProt protein accession numbers were provided by PEAKS upon peptide identification.

### Cellular component location embedding

Gene Ontology (GO) cellular component (CC) annotations for all UniProt protein IDs was obtained from uniprot.org. Pretrained 200 dimensional vectors for 64,649 GO terms were obtained from (Kim et al., 2021). Vectors were mapped to UniProt IDs and summed if a UniProt ID had more than 1 associated GO CC term. UMAP dimensionality reduction (McInnes et al., 2018) using the “hyperboloid” metric was applied, then mapped to the Poincare disk model. The resulting 2 values for each protein were used as features.

### IEDB Data

Peptides were selected from the Immune Epitope Database and Analysis Resource (www.iedb.org) (Vita et al., 2019) on July 16, 2021 using filters “Epitope Structure: Linear Sequence”, “Included Related Structures: neo-epitope”, “No B cell Assays”, “No MHC assays”, “MHC Restriction Type: Class I”, “Host: Homo sapiens (human)” and “Include Positive Assays”, “Include Negative Assays” (T cell assays). This resulted in 3754 peptides. Peptides whose “Assay Antigen Antigen Description” sequence did not match “Epitope Description” sequence were filtered, resulting in 3521 peptides. Peptides were also filtered out if they did not have an associated UniProt ID, found in the “Related Object Parent Protein IRI’’ field, resulting in 3367 peptides. Peptides were additionally removed if they were not in the UniProt CC file (downloaded on April 17, 2020 from uniprot.org by selecting all Human peptides and choosing “Gene ontology (cellular component)” in the column selection, see *Cellular component location embedding*), resulting in 3125 peptides. Peptides were dropped if they did not have a specific associated HLA allele (e.g. Allele Name = “HLA class I” or Allele Name = “HLA-A2”) or if they were not a simple linear sequence (e.g. ILCETCLIV + AIB(C3, C6)). This resulted in 2943 peptides. Affinity, stability, and foreignness features were calculated as described in “Validation Data”. Hex plots: Parent protein locations were plotted for each unique protein and immunogenic state for each study (i.e. if peptides from ProteinA had positive and negative tests, the parent protein would be retained twice, once in each immunogenic category).

### Random forest model

The RandomForestClassifier from sklearn v0.24.2 was used using random state 2021. The StratifiedKFold function was used to perform the 10-fold splits, also using random state 2021. The Youden indices for each fold were obtained by taking the threshold that had the greatest TPR-FPR (i.e. greatest area under the curve). The median Youden index was used to classify peptides as immunogenic or not for downstream analyses.

### Wells Data

Experimentally validated peptides were obtained from published supplementary tables S4 and S7 in (Wells et al., 2020). Peptides were mapped to parent proteins by iterating through all UniProt proteins and looking for a match to any peptide with a wildcard in the given mutated position (e.g. Python code to find the wildtype peptide corresponding to “FLCEILRSMSI” with mutated position 10: re.findall(r’(?=(“FLCEILRSM.I”))’, protein_sequence). 10 peptides without a mutated position were excluded. 584 (97.6%) of neopeptide sequences mapped to 1 unique parent wildtype sequence. Peptides with matched wildtype sequences mapping to multiple UniProt IDs were dropped. Missing foreginness or agretopicity scores were recalculated using the methods described in Wells *et al*. The resulting 558 peptides (Supplementary Table 2) from the discovery set was used to train a Random Forest classifier using sklearn (v0.24.2).

### Wells Validation Data

The trained model was tested on the 310 peptide validation dataset from Wells et al., as well as 43 peptides from ovarian tumors (Liu et al., 2019b). As these datasets did not include all features from the discovery dataset, NetMHCstabpan (v1.0) (Rasmussen et al., 2016) was used to predict pMHC stability, NetMHCpan (v4.0) (Jurtz et al., 2017) was used to predict pMHC binding affinity, and the antigen.garnish package (https://github.com/andrewrech/antigen.garnish) was used to calculate foreignness as described in Luksa, Wells. Finally, agretopicity was calculated by taking the ratio of mutant to wildtype binding affinity.

### Data and code availability

https://github.com/cartercompbio/Ploc

