## supplementary for "Immunogenic potential of neopeptides depends on parent protein subcellular location"

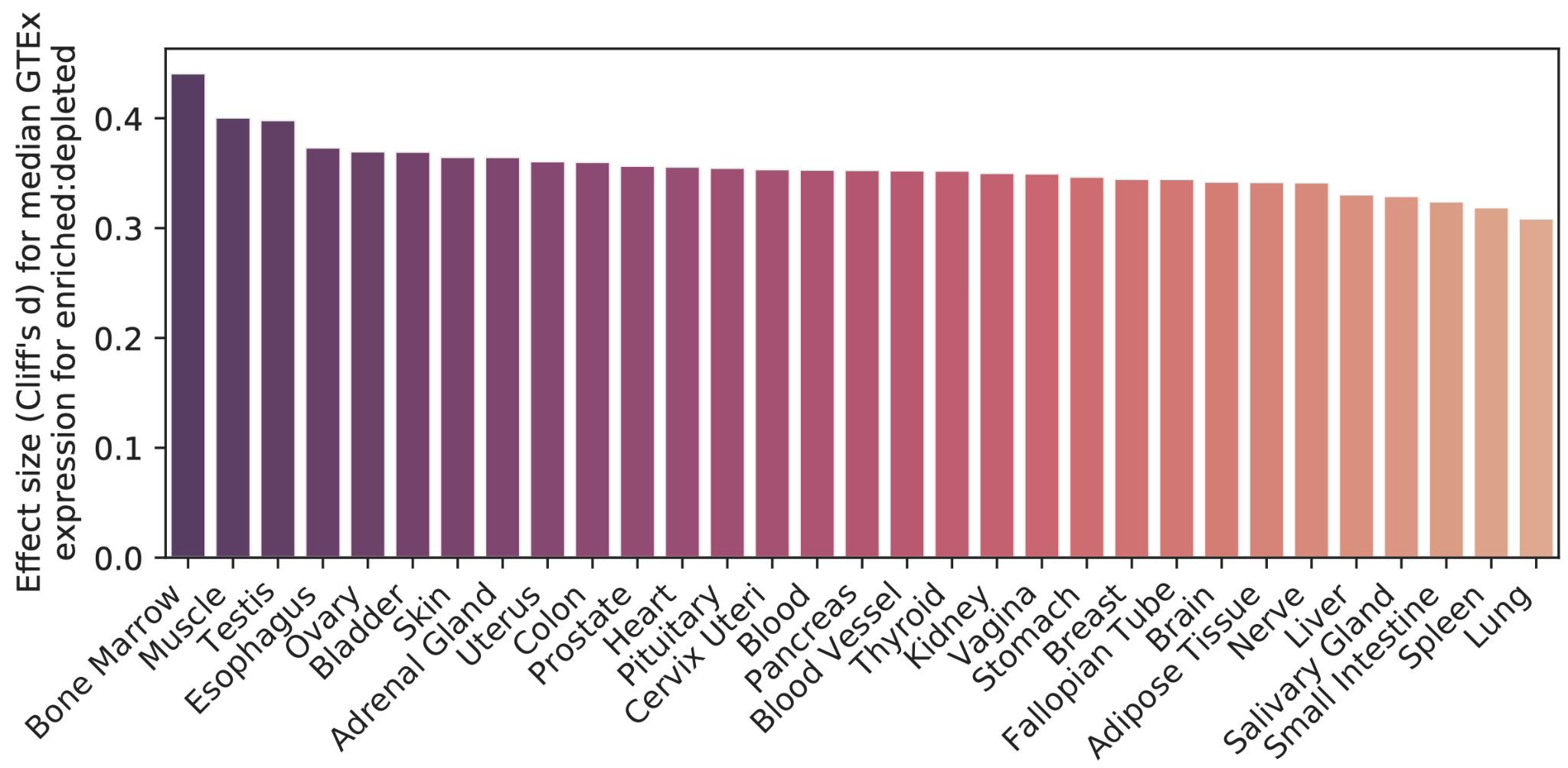

**Supplementary Figure 1**. Correlation of gene expression and eluted peptide location. Barplot for each tissue showing the effect size comparing the median GTEx gene expression for enriched over depleted genes. All comparisons show that genes in enriched locations have higher median gene expression than genes in depleted locations.

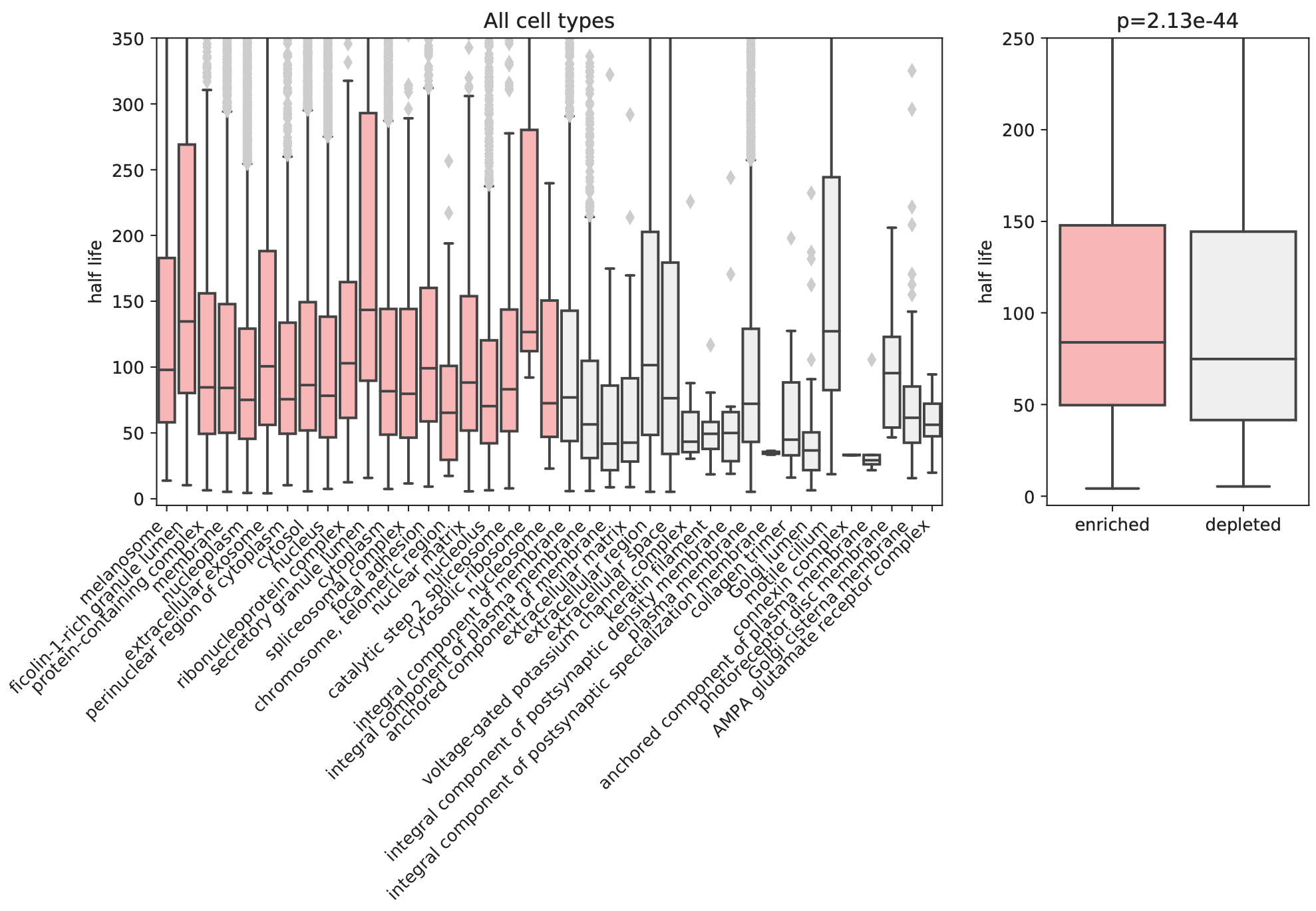

**Supplementary Figure 2**. Relationship between protein turnover and elution for the top 20 most frequently enriched or depleted cellular components across evaluated tissues or cell lines. The Mann-Whitney U test was used to compare statistical significance.

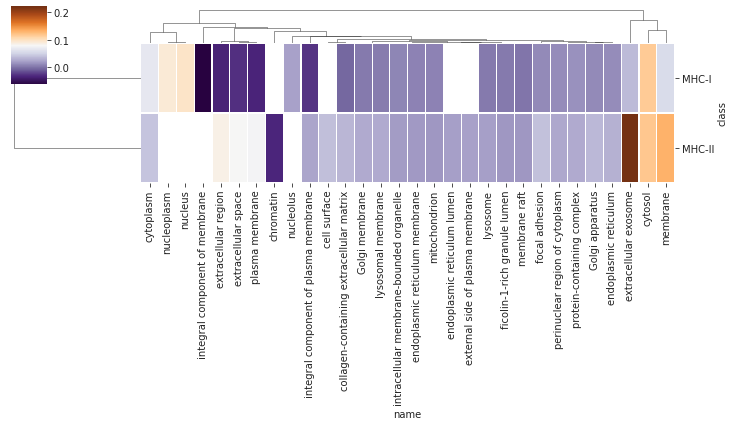

**Supplementary Figure 3**. Clustermap of significantly enriched (orange) or depleted (purple) cellular components for eluted pMHC class I and II from normal thymus. The color indicates the difference in study vs population enrichment.

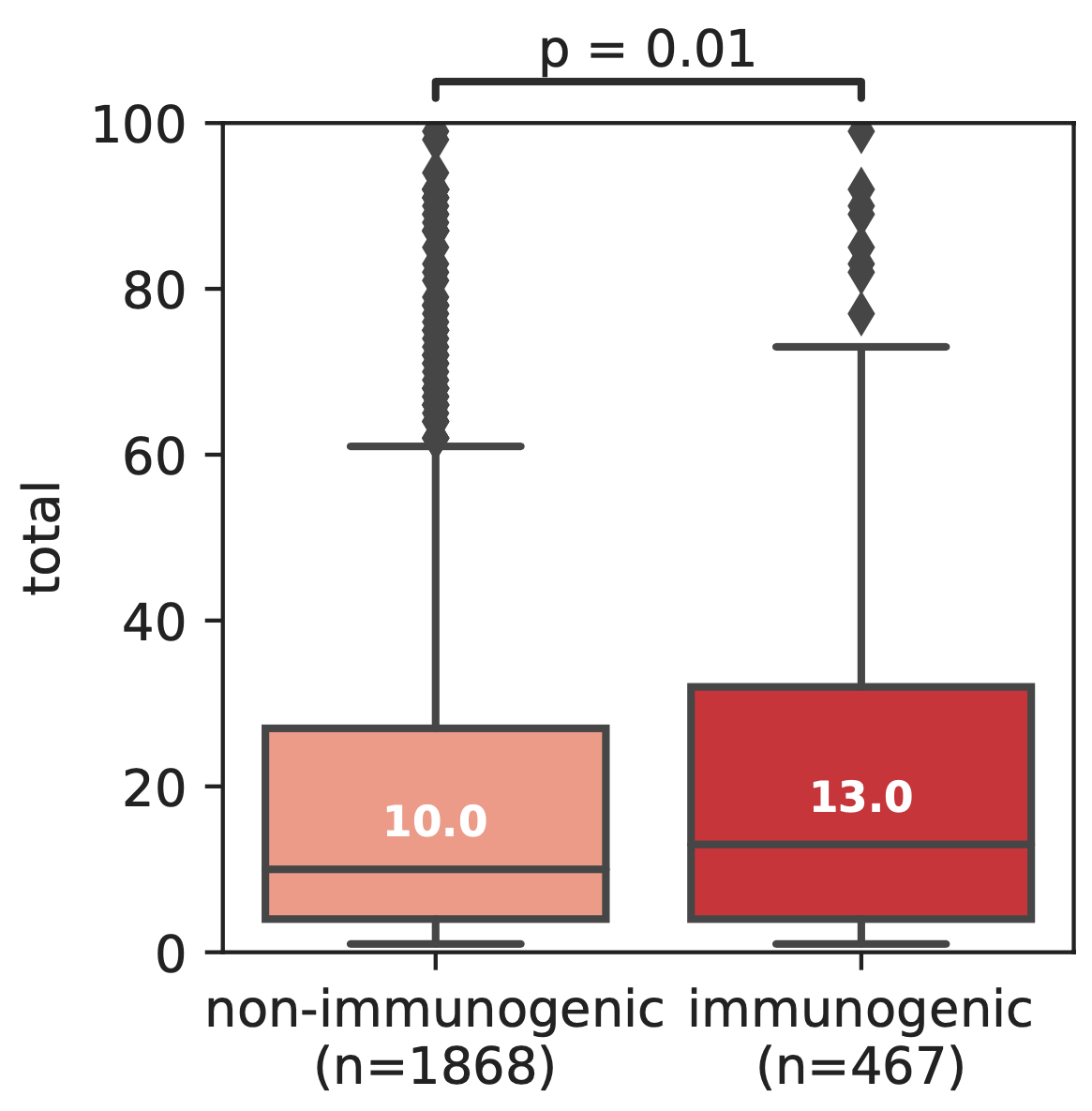

**Supplementary Figure 4**. Boxplot comparing the frequency of eluted peptides versus immunogenicity assay results for proteins that have been evaluated for class I immunogenicity in the IEDB.

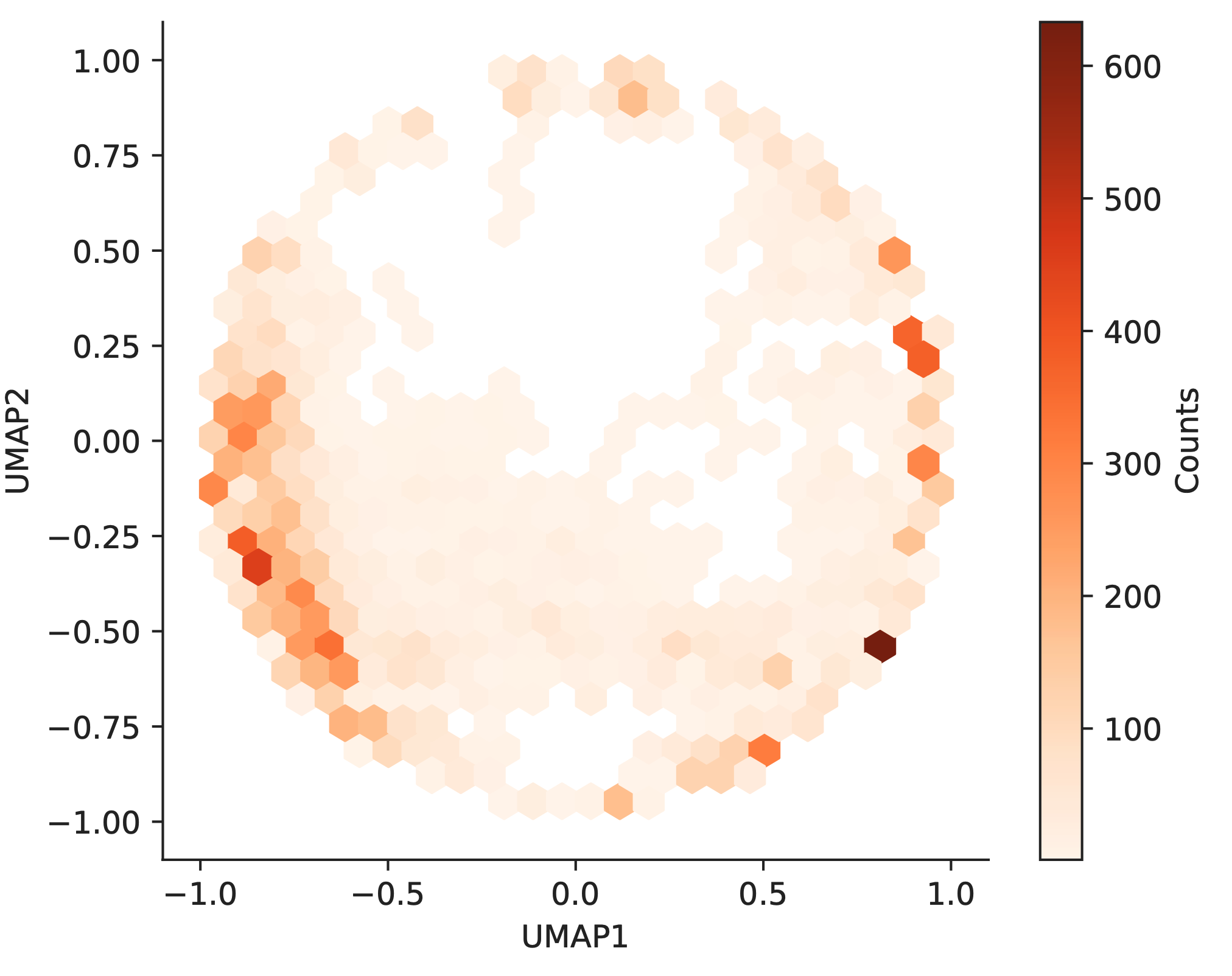

**Supplementary Figure 5**. Hexplot of UMAP location embeddings for all unique UniProt proteins with reviewed status and unique gene names.

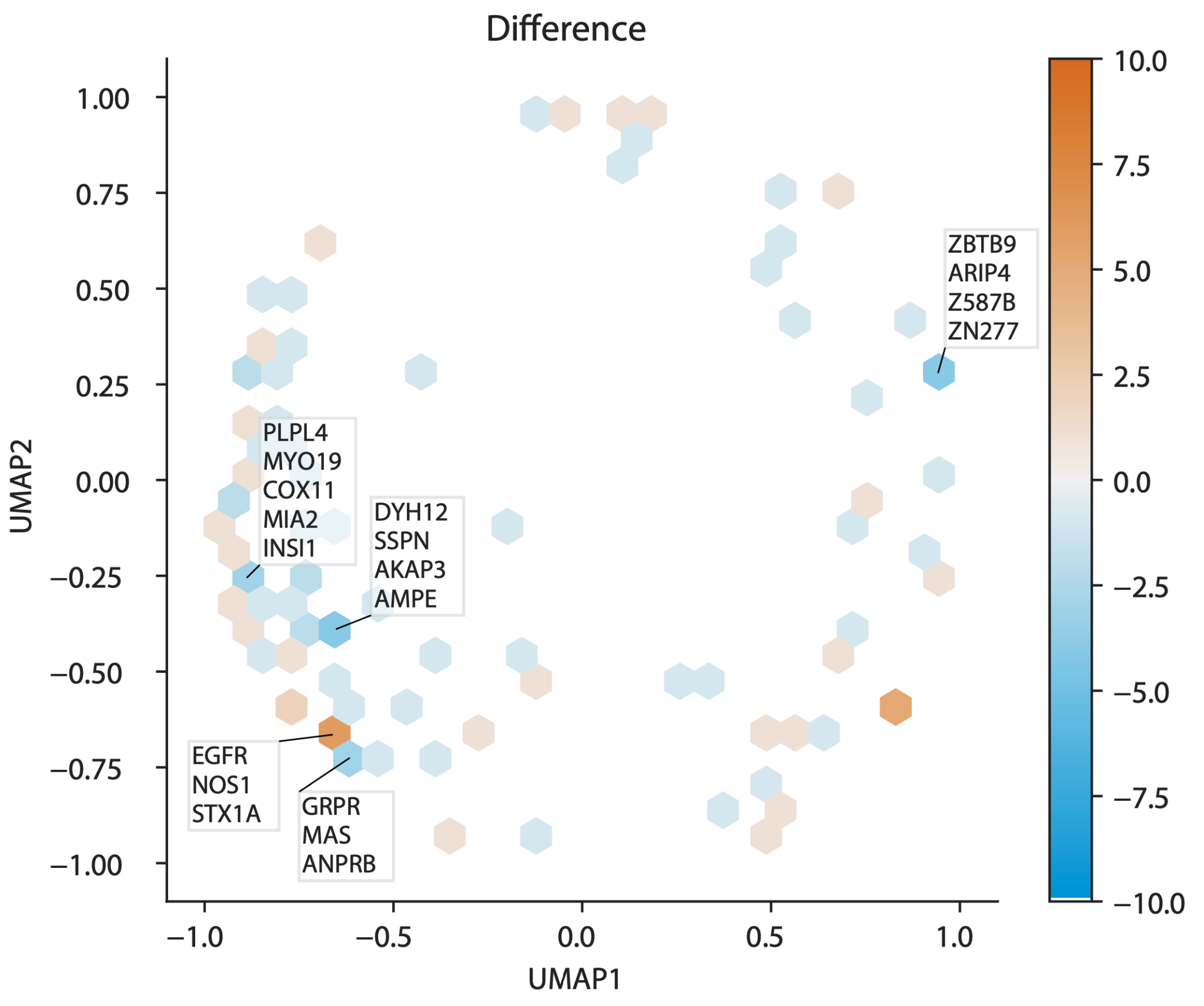

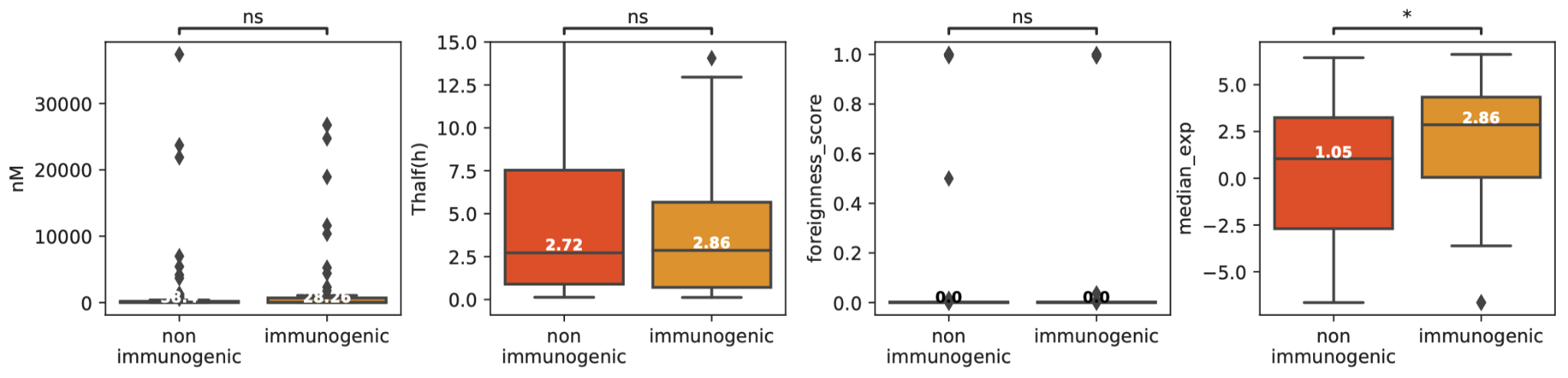
**Supplementary Figure 6**. Overview of differentially classified peptides between the models with and without location as a feature. Orange indicates locations with more immunogenic peptides compared to non-immunogenic peptides and vice versa. Locations with more than 3 immunogenic or non-immunogenic genes are highlighted.

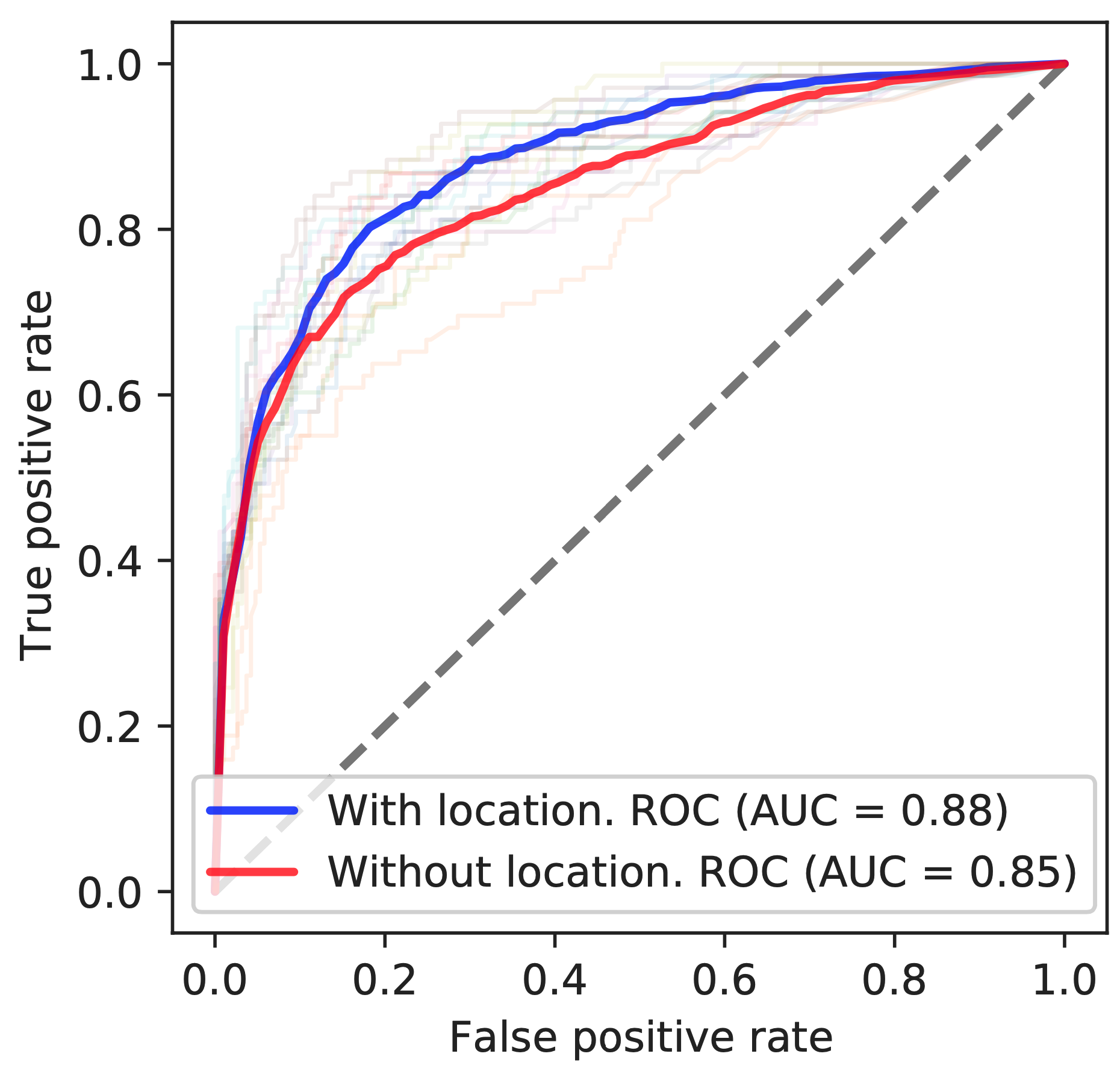

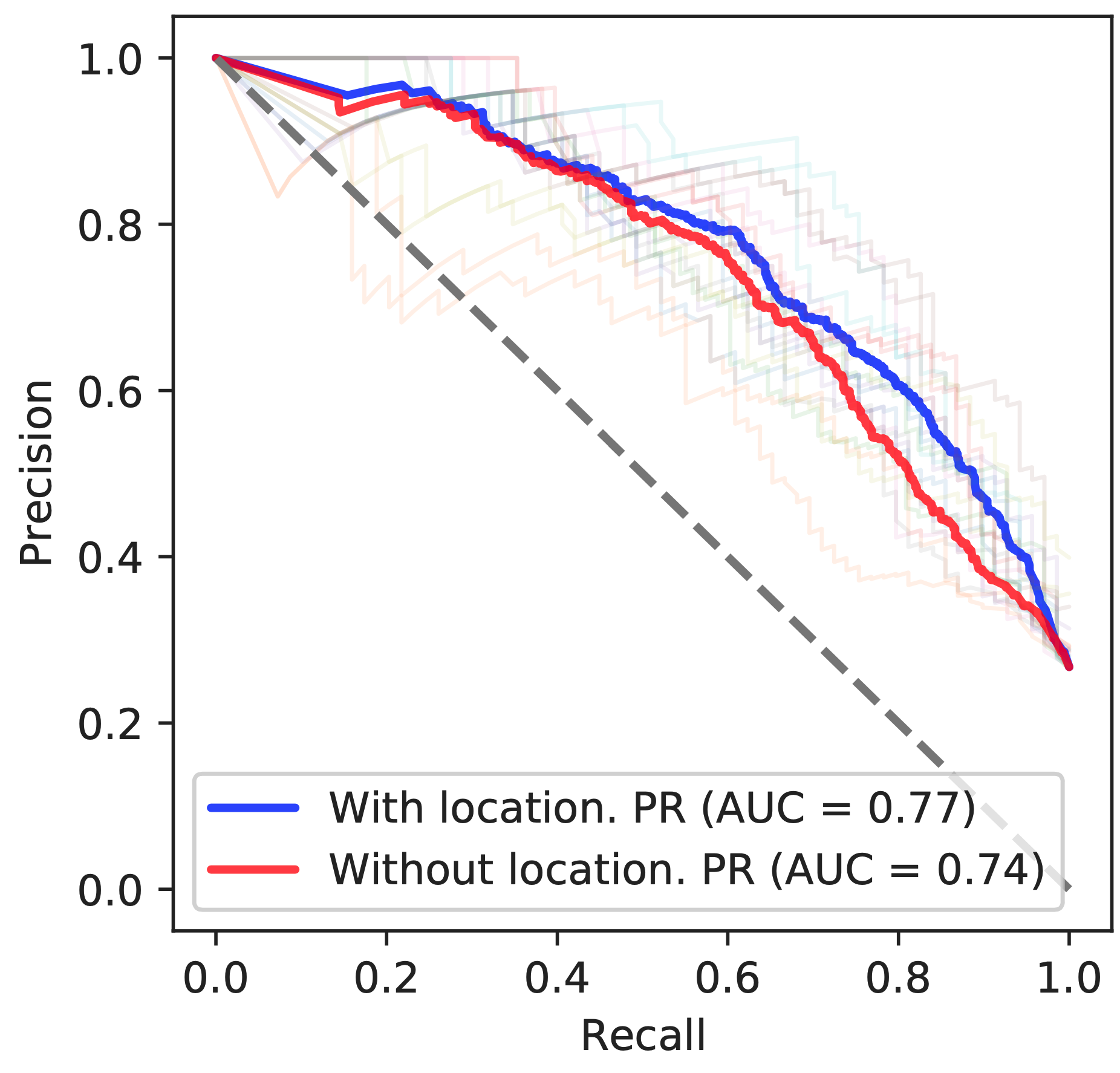

**Supplementary Figure 7**. (Left) Area under the receiver operating characteristic curve (AUROC) and (right) area under the precision recall curve (AUPRC) for 10-fold cross validation using a Random Forest model incorporating median GTEx gene expression, peptide affinity, stability, and foreignness (Methods) with and without parent protein location features.

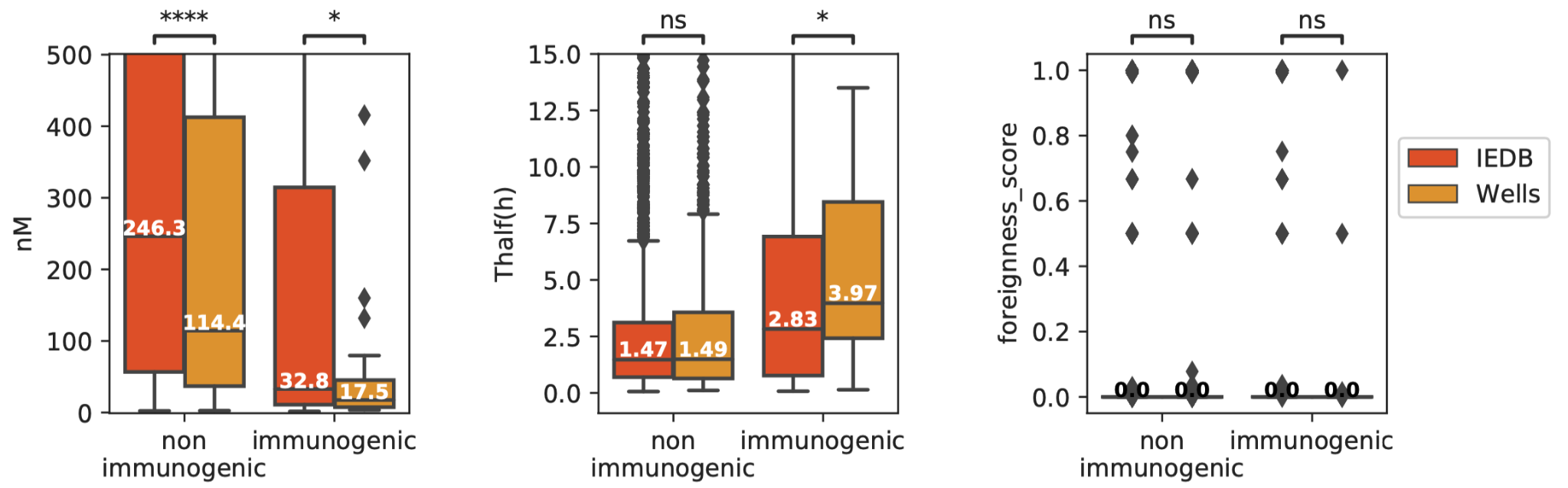

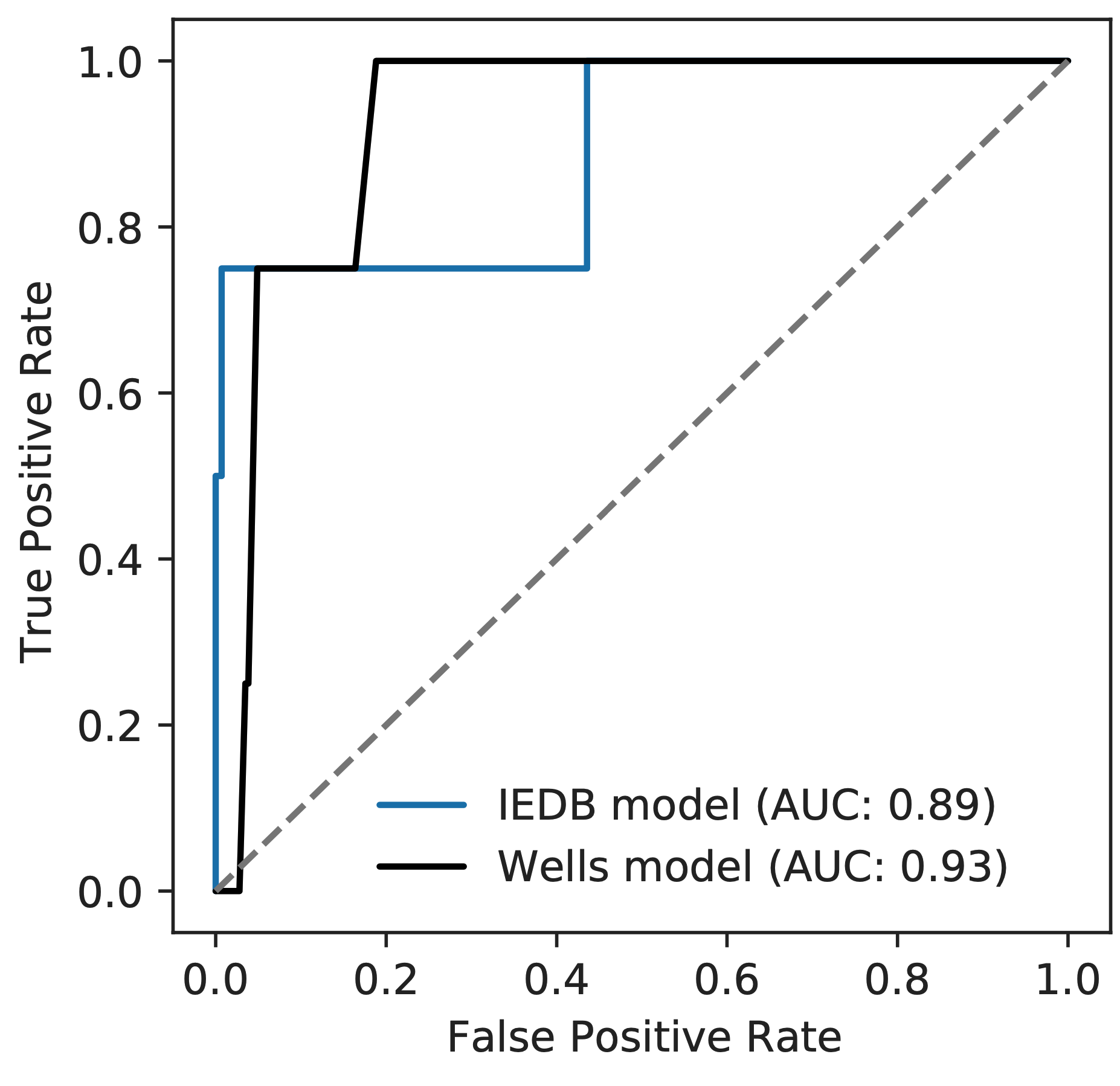

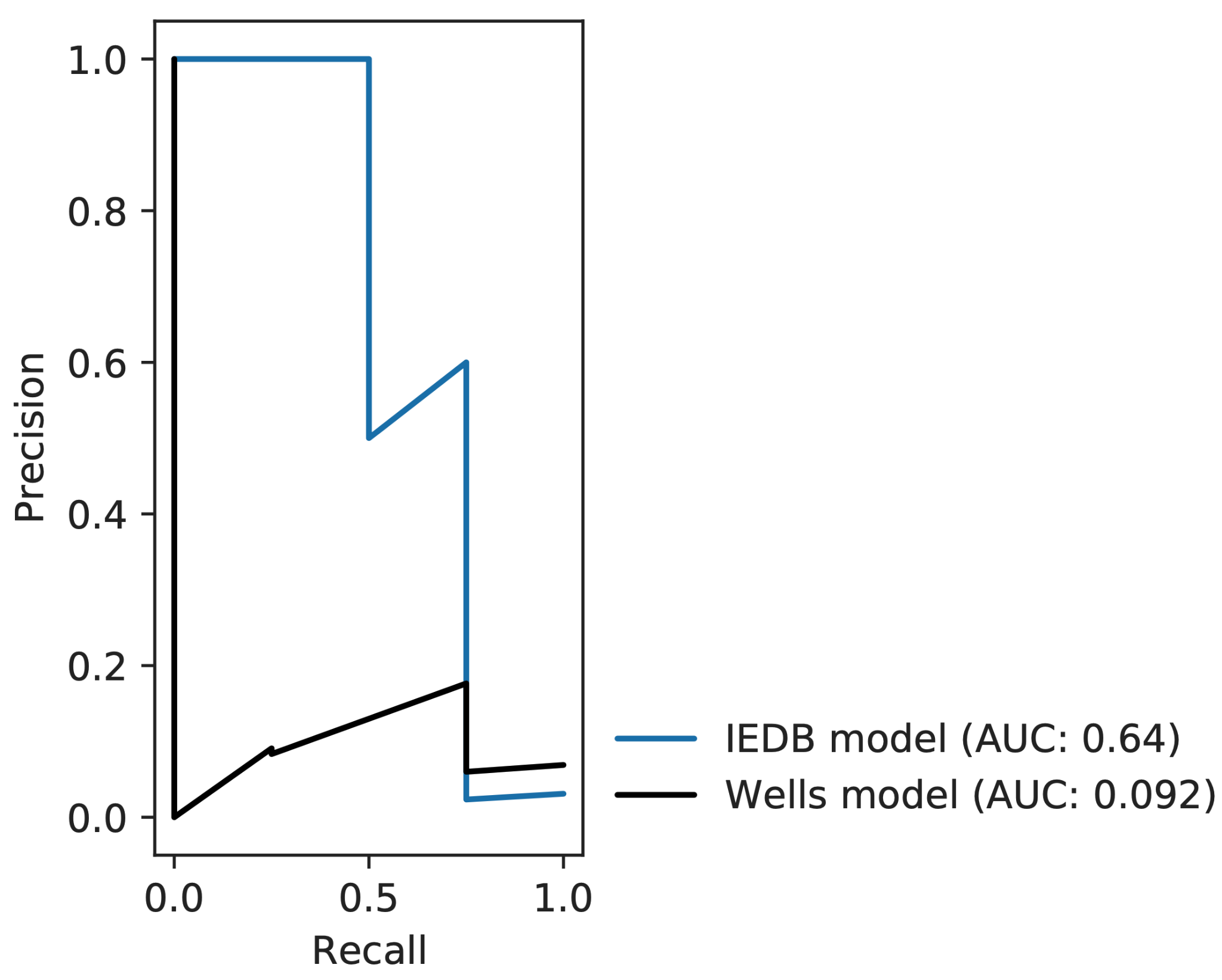

**Supplementary Figure 8**. Comparison of the IEDB and Wells *et al*. datasets. (Top) Boxplots comparing affinity (measured in nM), stability (measured by half life), and foreignness stratified by immunogenicity. The Mann-Whitney U test was used to compare statistical significance. (Bottom panel) Area under the receiver operating characteristic and precision recall curves using the random forest model trained on the IEDB dataset and Wells discovery set to test on the Wells test set.

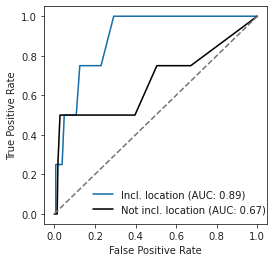

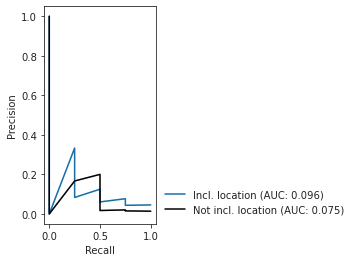

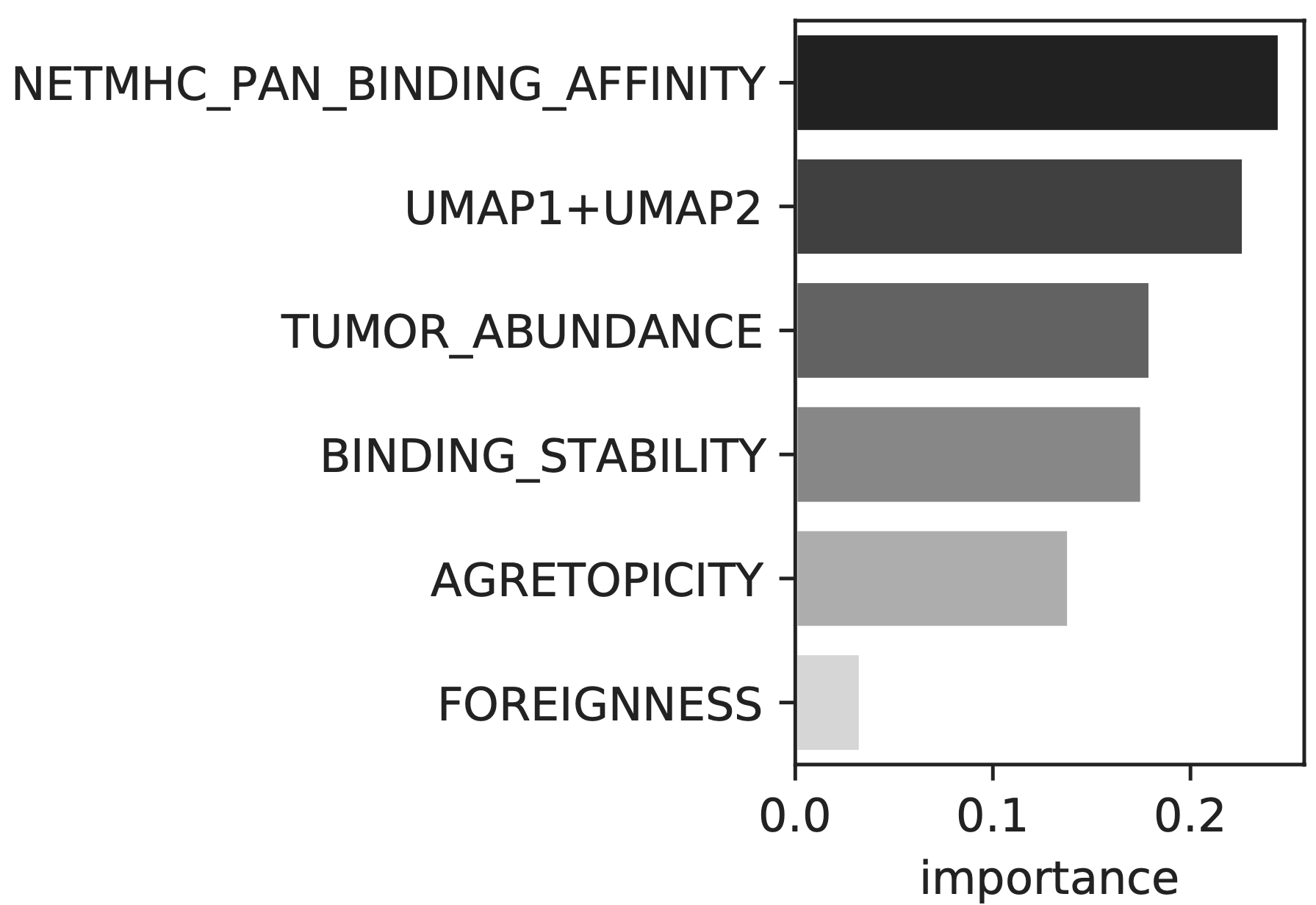

**Supplementary Figure 9**. AUROC and AUPRC plots for the model trained on the Wells discovery dataset and tested on the Wells test dataset. Features include peptide affinity, stability, tumor abundance, agretopicity, and foreignness. Barplot denoting feature importance for the model.

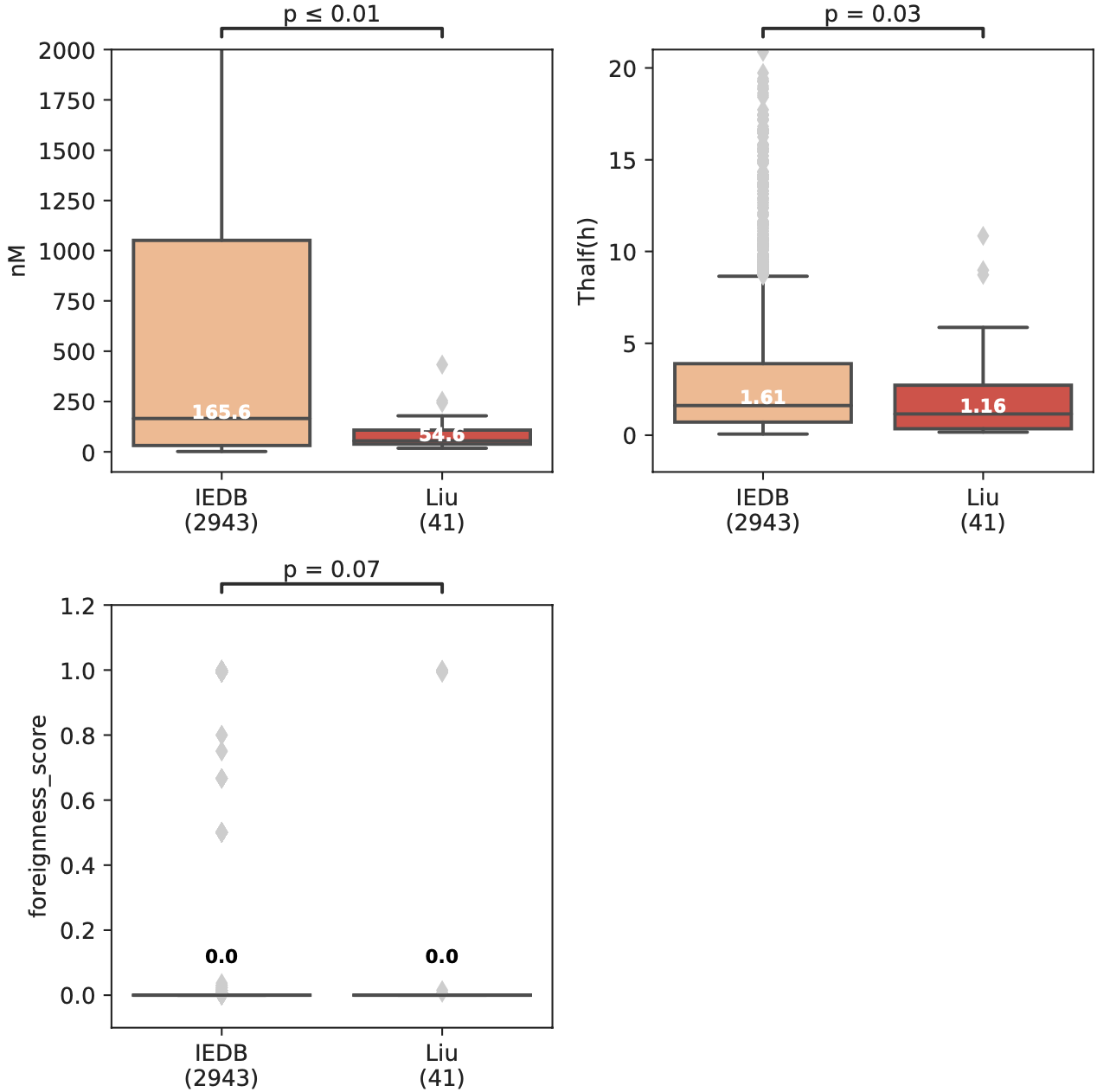

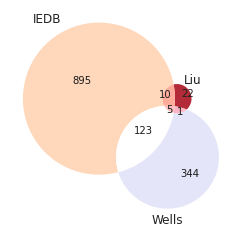

**Supplementary Figure 10**. Comparison of the IEDB and Liu *et al*. datasets. Boxplots comparing affinity (measured in nM), stability (measured by half life), and foreignness. The Mann-Whitney U test was used to compare statistical significance. The Venn diagram shows the overlapping unique locations. All overlapping locations were non-immunogenic in Liu.

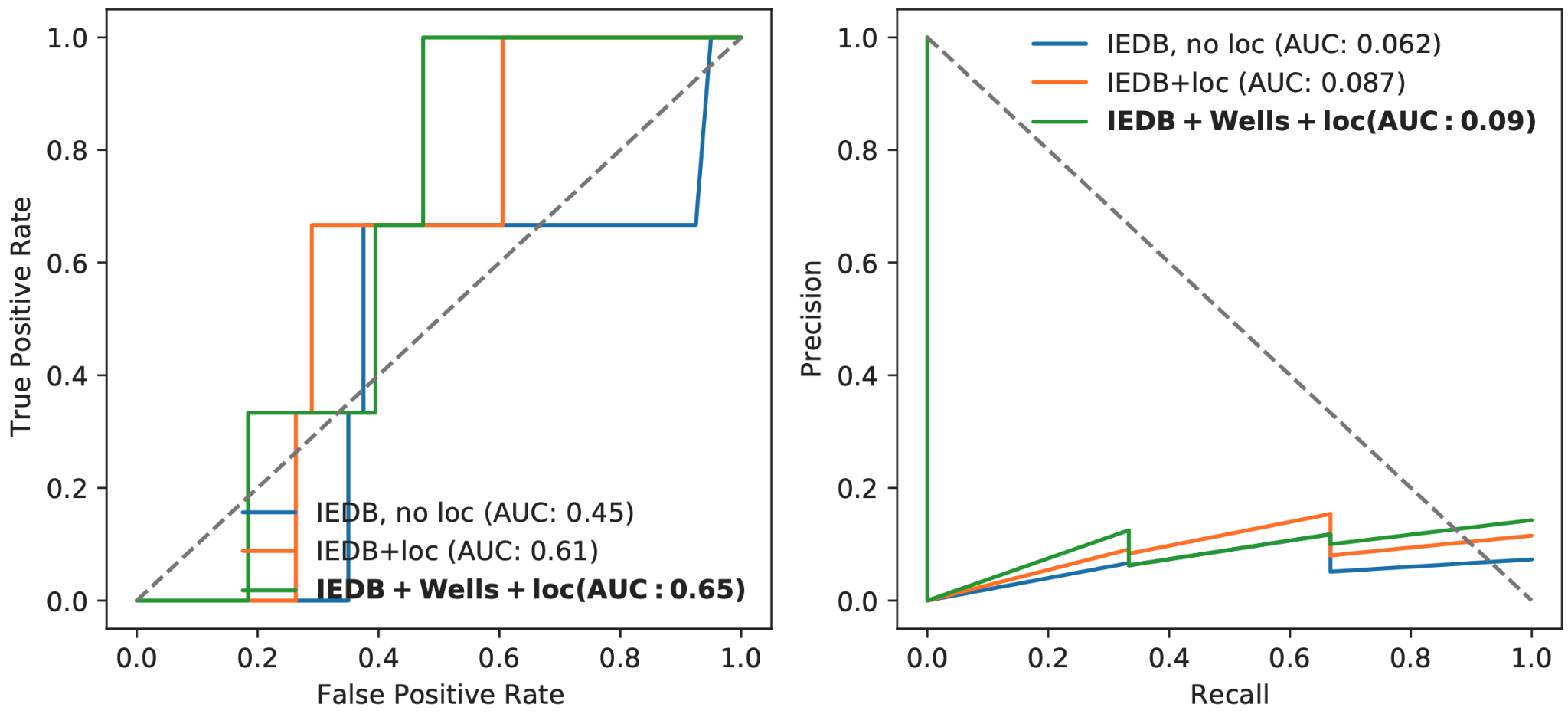

**Supplementary Figure 11**. Testing pretrained models on the unseen Liu ovarian dataset. (Left) AUROC and (right) AUPRC curves for the IEDB model without location, with location, and aggregated model with IEDB, Wells et al., and location.

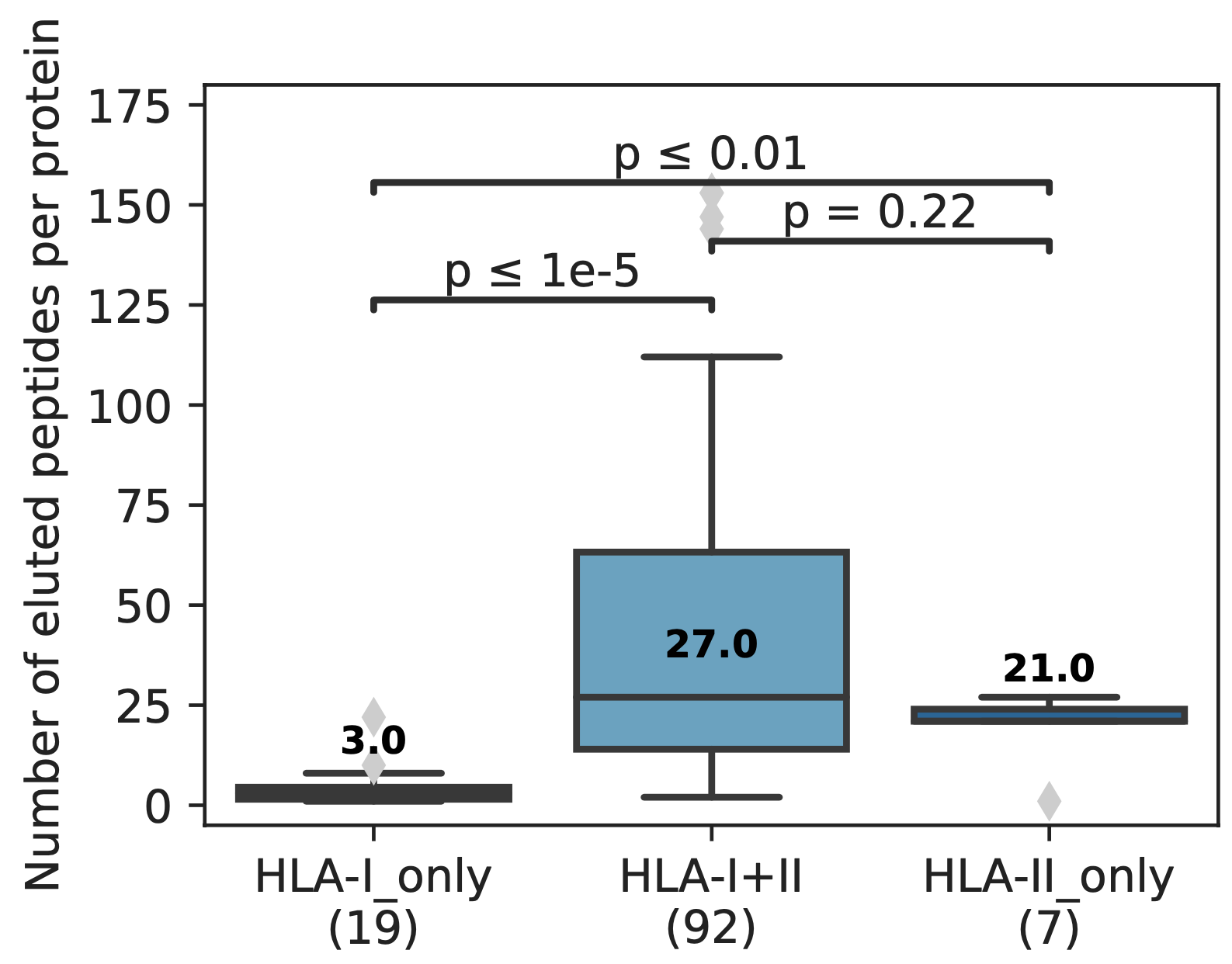

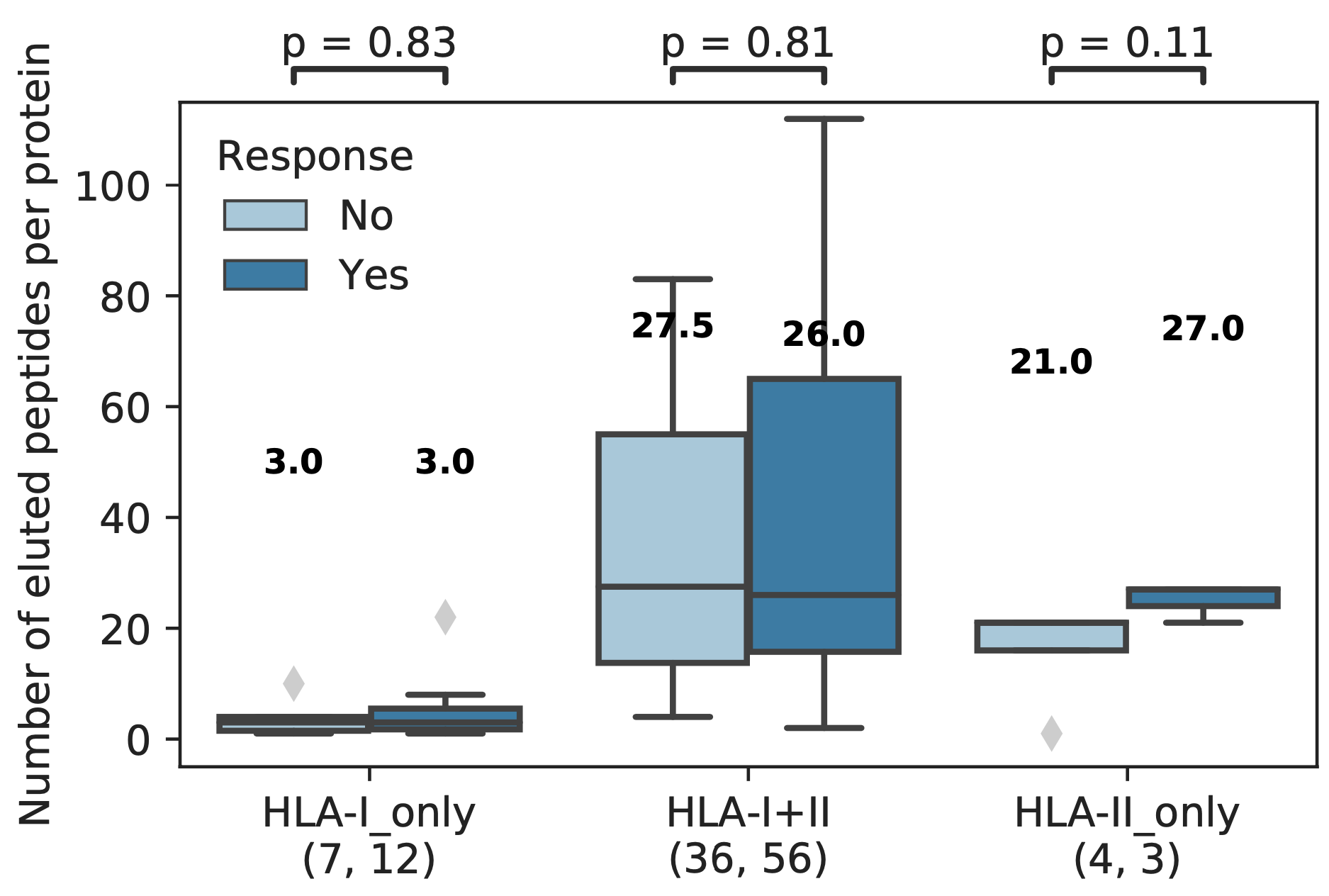

**Supplementary Figure 12**. Analysis of neopeptide vaccine parent protein MHC elution patterns. A) Boxplots showing the number of eluted peptides in the HLA ligand atlas associated with the parent proteins of the 125 neopeptides evaluated by Sahin *et al*. The majority were from proteins from which peptides were found in both MHC-I and MHC-II eluted complexes. Parent proteins exclusive to MHC-I tended to have lower eluted peptide counts than parent proteins exclusive to MHC-II. B) Boxplots showing the number of eluted peptides as in panel A, but further divided according to whether the The number of MHC eluted peptides was not associated with post-vaccination response.

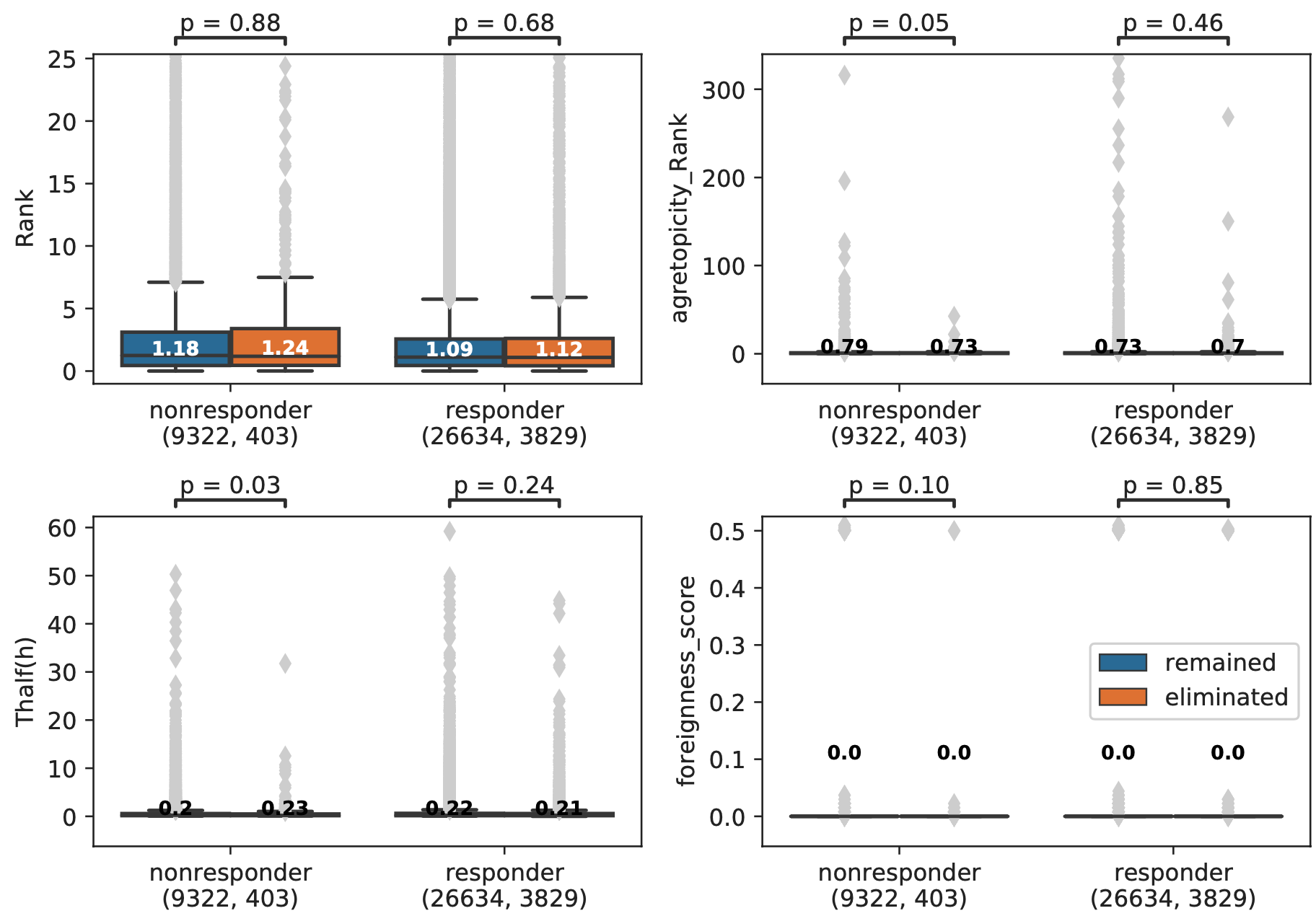

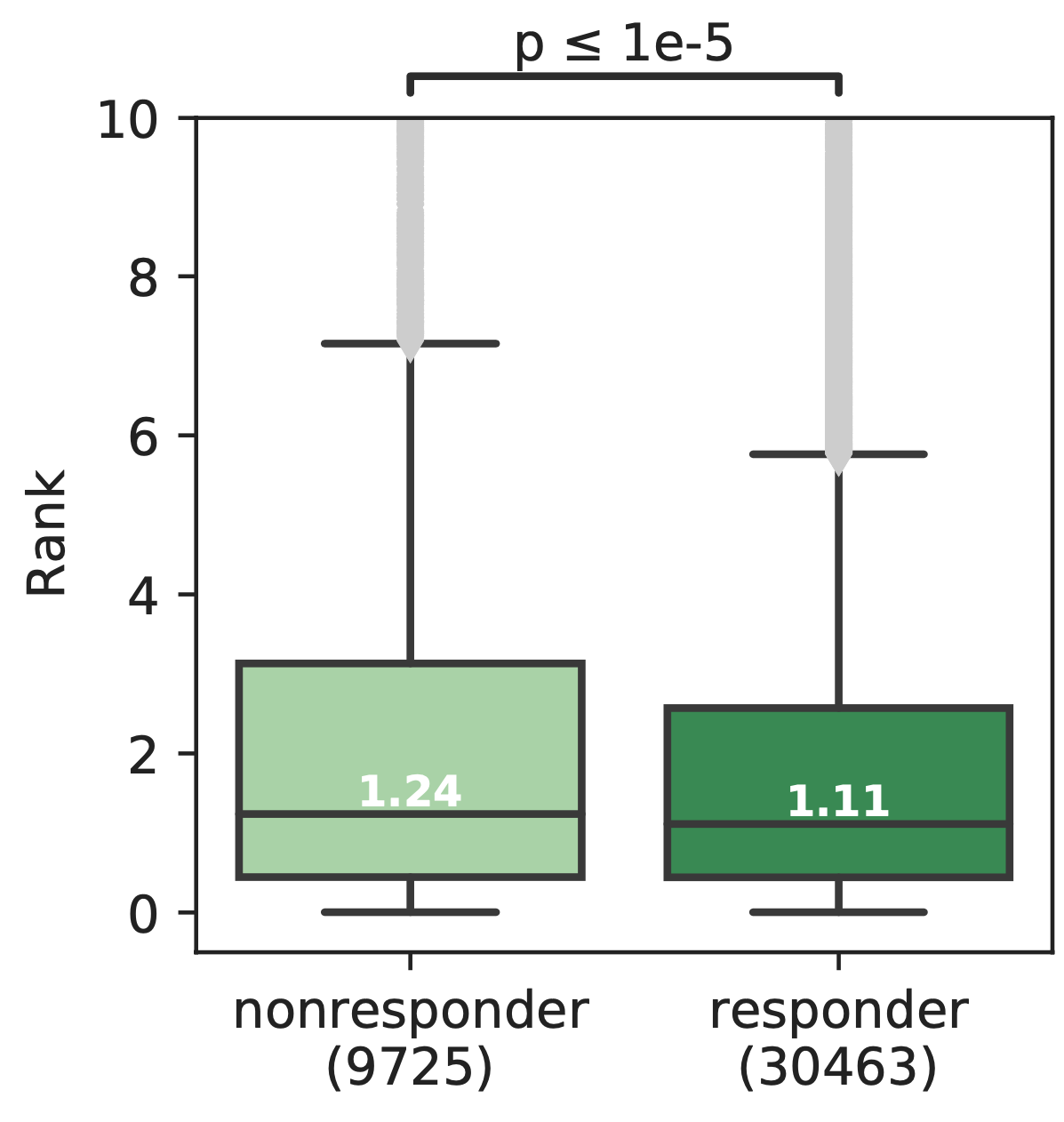

**Supplementary Figure 13**. Comparison of neopeptide characteristics in the Riaz *et al*. dataset. (Top) Boxplots comparing affinity, agretopicity, stability, and foreignness between eliminated versus remaining neopeptides for both responders and nonresponders. (Bottom) Comparison of neopeptide affinity between responders and nonresponders.

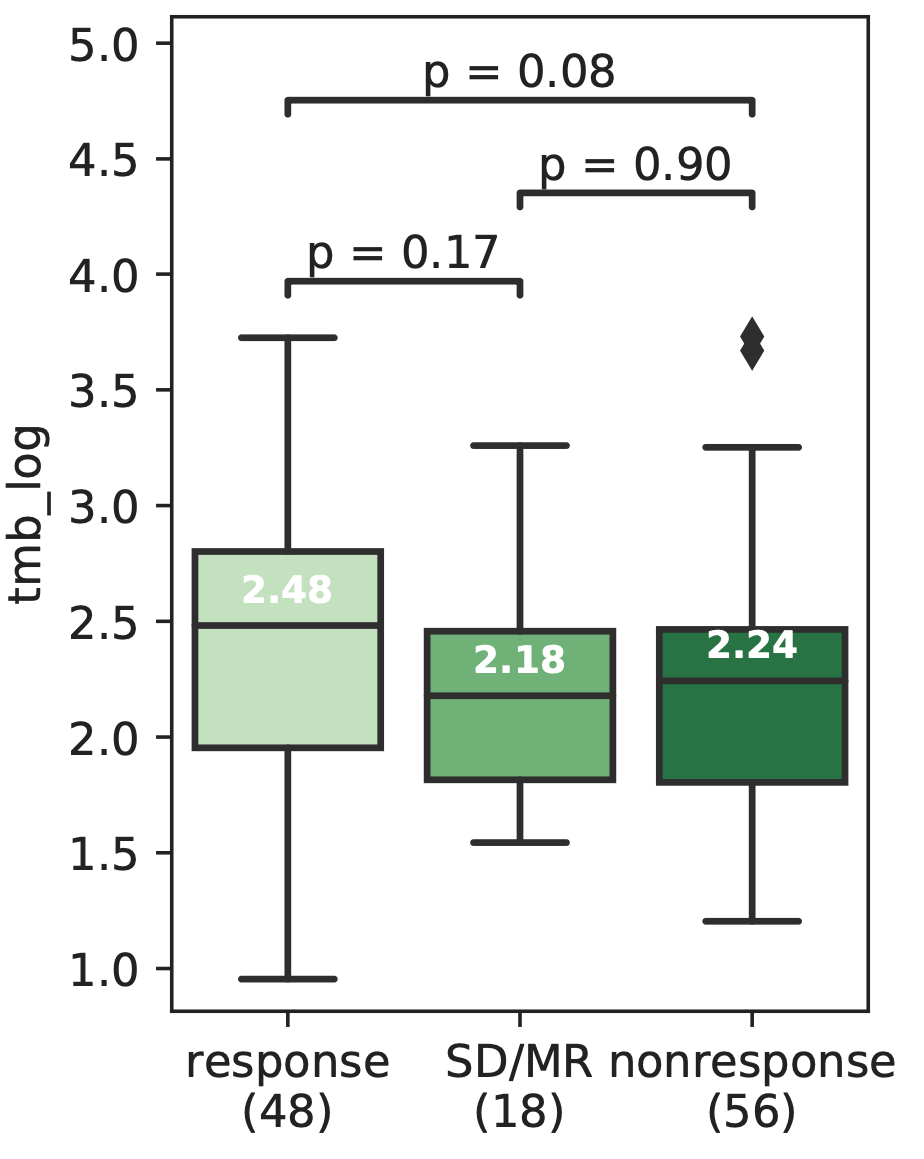

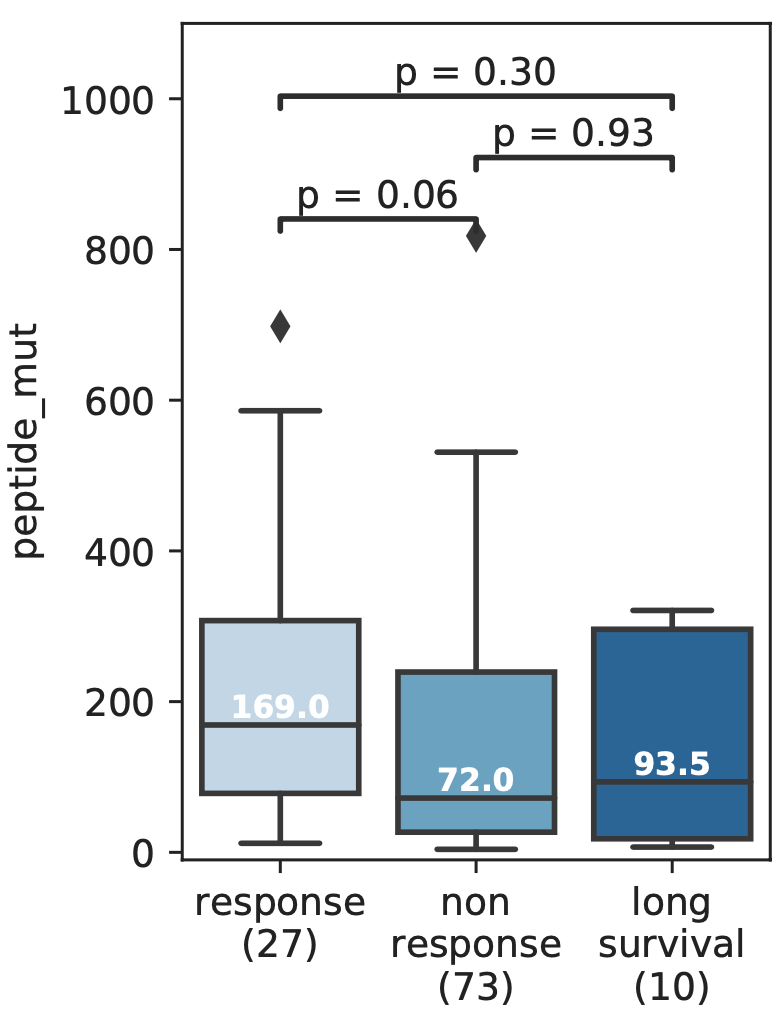

**Supplementary Figure 14**. Initial association of tumor mutation burden with response in (A) Liu *et al.* and (B) Van Allen *et al.* Tumor mutation burden was defined in Liu *et al*. as the number of somatic mutations, and in Van Allen *et al.* as the number of neopeptides <500nM.

**Supplementary Table 1.** Cox proportional hazards results

|  |  | **Hazard ratios (exp(coef))** | **Standard error (se(coeff))** | **p-value** |
| --- | --- | --- | --- | --- |
| Model focusing on the 40 genes that fall in locations seen to be previously immunogenic  Partial AIC=401.7 | PHBR score | 1.29 | 0.21 | 0.24 |
|  | TMB | 0.99 | 0.004 | 0.15 |
|  | Age at treatment with immunotherapy | 1.00 | 0.012 | 0.97 |
|  | Gender | 1.16 | 0.30 | 0.62 |
| Unfiltered model. Focuses on all genes from the gene panel  Partial AIC=400.6 | PHBR score | 1.11 | 0.057 | 0.069 |
|  | TMB | 0.99 | 0.004 | 0.13 |
|  | Age at treatment with immunotherapy | 1.00 | 0.01 | 0.94 |
|  | Gender | 1.18 | 0.28 | 0.58 |
